# Discovery and Biosynthesis of Nitrilobacillins by Post-translational Introduction of C-Terminal Nitrile Groups

**DOI:** 10.64898/2026.03.11.711119

**Authors:** Lide Cha, Chuyang Qian, Chandrashekhar Padhi, Lingyang Zhu, Wilfred A. van der Donk

## Abstract

Nitrile-containing natural products are produced in all kingdoms of life. Despite the wide application of nitrile-containing peptide scaffolds in medicinal chemistry, the presence of the nitrile group is unprecedented in ribosomally synthesized and post-translationally modified peptides (RiPPs). In this work, we report the identification and characterization of a RiPP biosynthetic gene cluster (BGC), where an asparagine synthetase-like (AS-like) protein encoded in the BGC converts the C-terminal carboxylate of the precursor peptide to a nitrile. Furthermore, a multinuclear nonheme iron-dependent oxidative enzyme (MNIO) and an α-ketoglutarate-dependent HExxH motif-containing enzyme (αKG-HExxH) perform stereoselective β-hydroxylation of aspartate and proline residues, respectively. Structure prediction-guided mechanistic evaluation of the nitrile synthetase provided insights into the possible mechanism of catalysis. These findings extend our understanding of the structural diversity of RiPPs and demonstrate the catalytic versatility of AS-like enzymes in natural product biosynthesis.

## Introduction

Since the discovery of allyl cyanide in 1863,^1^ nitrile-containing natural products have been reported to be produced by animals, plants and microorganisms.^2–5^ The electrophilic nature of the nitrile carbon contributes to the activity of nitrile-containing metabolites in biology and medicinal chemistry.^6–8^ In nature, production of the nitrile group is achieved mostly via dehydration of the corresponding aldoximes catalyzed by various enzymes (Figure S1a).^9,10^ Alternatively, nitrile formation proceeds through oxidative or ATP-dependent mechanisms (Figure S1a).^5,11–17^ Nitrile groups have been identified in diverse classes of natural products including glycosides, alkaloids, and terpenes, but nitrile-containing peptide natural products are rare, with auranthine the only example discovered thus far (Figure S1b). In contrast, use of the nitrile functional group is common in synthetic peptides designed to inhibit proteases, with one notable example being nirmatrelvir, a key ingredient in the SARS-CoV-2 therapeutic Paxlovid that inhibits the viral main protease (Figure S1b).^18^

Recent estimates suggest that a very large fraction of natural products (up to 97%) remains to be discovered.^19^ Peptide secondary metabolites are mostly derived from non-ribosomal or ribosomal pathways.^20–22^ Compared with non-ribosomal pathways, which include the biosynthesis of auranthine, ribosomally synthesized and post-translationally modified peptides (RiPPs) are differentiated by precursor peptides that are genetically encoded. Estimations of the abundance of various natural product classes in defined natural environments show that RiPPs are a major family,^23^ and in some environments such as the human microbiome, RiPPs appear to be the most prevalent class of natural products.^24^ Maturation of RiPPs involves diverse post-translational modifications (PTMs) of the precursor peptide.^22^ Around 50 classes of RiPPs have been reported to date that are categorized by class-defining PTMs. The number of distinct PTMs continues to expand as a result of the rapid increase in genome sequences over the past two decades.^25^ A large number of different enzyme families have been linked to RiPP biosynthesis.^26,27^ Because the substrate sequences of RiPP modifying enzymes are encoded within the BGC, functional characterization of RiPP biosynthetic enzymes for which activity has not been reported previously is less challenging compared with their counterparts in the biosynthesis of other natural products. These features have made RiPP BGCs excellent repositories to discover novel enzyme chemistries.^25^

Recent studies have expanded the chemical space of the products of non-heme iron dependent enzymes in RiPP biosynthesis,^28–35^ including RiPP BGCs that encode MNIO and αKG-HExxH proteins. In the current work, we focused on a BGC that contains members of both enzyme classes from *Peribacillus simplex* VanAntwerpen02. We demonstrate that the MNIO and HExxH enzymes catalyze β-hydroxylation of aspartate and proline residues, respectively. The more unique reaction in the pathway features an AS-like enzyme that unexpectedly installs a nitrile group at the C-terminus of the precursor peptide. Mechanistic investigation of this enzyme suggests that the nitrilation reactivity emerged from the canonical asparagine synthetase catalytic machinery. We termed the products from this BGC nitrilobacillins because orthologous pathways were only identified in *Bacillus*-related genera. Production of the nitrile-containing peptide seems to be regulated by two different mechanisms. First, nitrile installation requires Asp hydroxylation by the MNIO. Second, an apparent pseudo enzyme related to the MNIO protein is encoded within the BGC, which can compete with the MNIO protein in binding to the MNIO partner protein, thereby preventing nitrile biosynthesis. This work expands RiPP biosynthesis with a pharmaceutically significant modification.

## Results

### Identification of the *pes* BGC

During our genome mining efforts for novel RiPP BGCs encoding MNIOs, a BGC (*pes* cluster) from *Peribacillus simplex* VanAntwerpen02 was identified. The BGC encodes the αKG-HExxH protein PesO, two precursor peptides PesA1/A2, an MNIO enzyme PesH, a putative MNIO partner protein PesI, an AS-like enzyme PesC, a hypothetical protein PesX, two M16 peptidases PesP1/P2, and a transporter protein PesT (Figure 1a). Unlike most previously reported MNIO enzymes, which require a partner protein that is encoded adjacently within the BGC,^28,30,33,36–39^ the putative MNIO partner protein gene *pesI* is positioned next to *pesX*. Initial bioinformatic analysis of PesX suggested that it is a hypothetical protein from an unknown protein family, but structural analysis with AlphaFold 3^40^ as well as a Foldseek^41^ search revealed that PesX likely possesses a triose-phosphate isomerase (TIM)-barrel fold like MNIOs and shares structural similarity to another MNIO protein MbnB (Figure S2). However, a sequence alignment of PesX with characterized MNIO proteins demonstrated that it lacks the iron-binding residues that are conserved in MNIOs (Figure S3).^42^ Therefore, the role and function of PesX was unclear.

**Figure 1.**
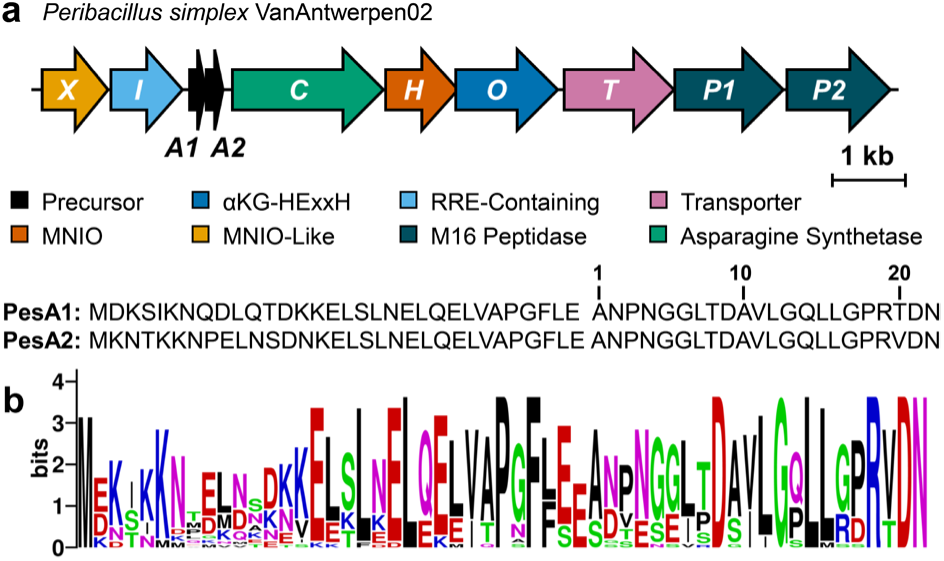
**a)** Composition of the *pes* BGC from *Peribacillus simplex* VanAntwerpen02 and the amino acid sequence of precursor peptides PesA1/2. The numbering is based on the C-terminal fragment obtained upon GluC-digestion; **b)** Sequence logo of precursor peptides from BGCs homologous to the *pes* BGC involving 42 total sequences and 21 unique sequences (Figure S5, identical sequences were removed).

A sequence homology search of the enzymes in the *pes* cluster using BLASTp, followed by genome neighborhood analysis using RODEO^43^ lead to the discovery of 26 gene clusters that resemble the *pes* cluster composition. These orthologous clusters all contain homologs of PesX, PesI, PesH and PesC. αKG-HExxH proteins are only present in ∼75% of the clusters and when they are absent, they are replaced by a predicted arginase (e.g. Figure S4). Alignment of the precursor peptides from these homologous clusters reveals several conserved residues including a conserved C-terminal R(V/T)DN motif (Figure 1b). Notably, a proline residue before the R(V/T)DN motif only co-occurs when an αKG-HExxH protein is encoded in the BGC, hinting that the latter enzyme may modify this Pro residue.

### Characterization of the product of the *pes* BGC

Because the strains encoding the native *pes* BGC were not available to us, we investigated the functions of the encoded enzymes by heterologous expression in *E*. *coli*. A His_6_-tag was appended to the N-terminus of the precursor peptides, and they were expressed separately or co-expressed with a subset of putative modifying enzymes. Following immobilized metal affinity chromatography (IMAC) purification, the purified peptides were digested with endoproteinase GluC and characterized by mass spectrometry.

We co-expressed both precursor peptides PesA1 and PesA2 individually with the various enzymes encoded in the *pes* BGC with essentially the same results. Considering the putative core region of PesA1 and PesA2 differs by only one residue (Figure 1a), we will describe the data with PesA2 here and refer to the Supporting Information for the analogous data for PesA1. Co-expression of PesA2 with the MNIO protein PesH and its potential partner PesI and analysis by liquid chromatography-high resolution mass spectrometry (LC-HRMS) revealed the emergence of a new species with a +16 Da mass shift with respect to the unmodified PesA2 (Figure 2). The site of PesHI modification was located to Asp21 by high-resolution tandem mass spectrometry (HR-MS/MS) (Figure S6).

**Figure 2.**
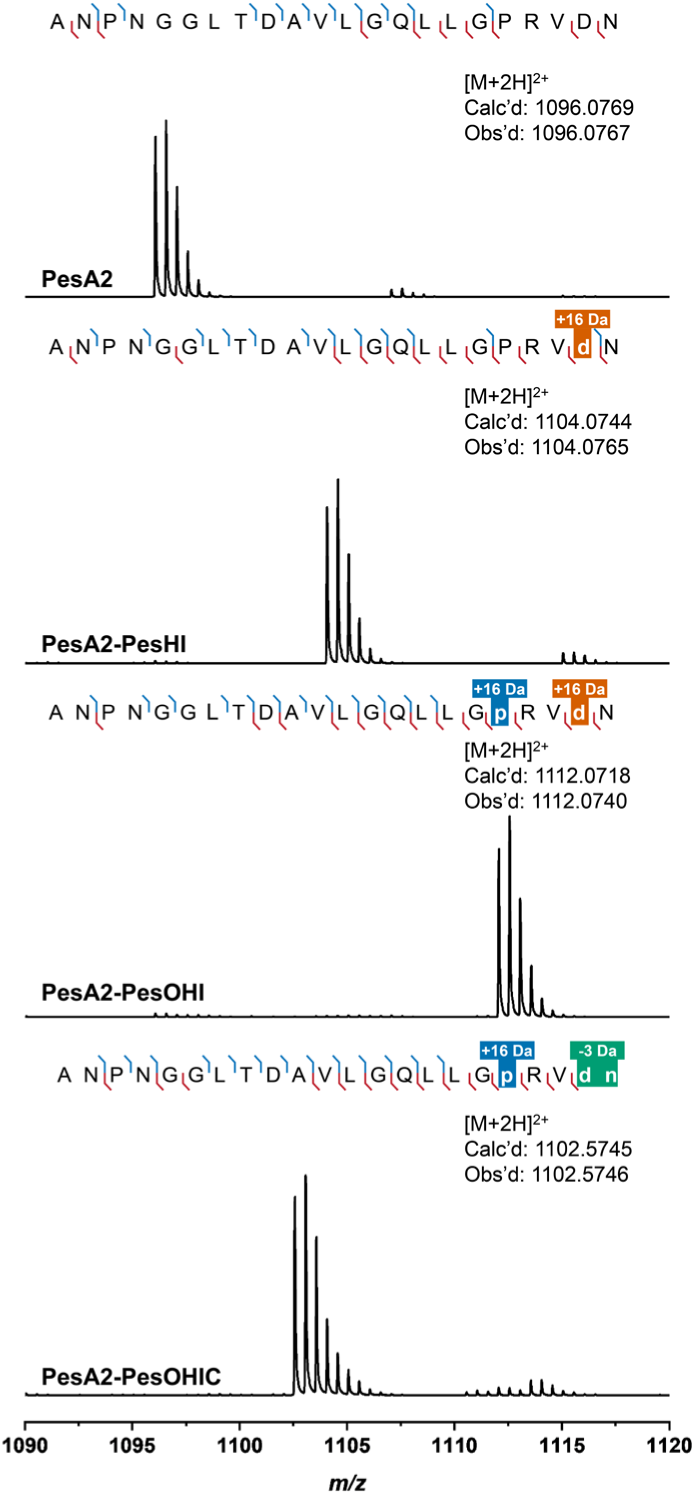
HR-MS/MS analysis of unmodified precursor peptide PesA2 or PesA2 co-expressed with the indicated modifying enzymes. The MS/MS fragmentation pattern of each modified peptide is shown (for tandem MS spectra, see Figure S6). Prior to analysis, the peptides were digested with endoproteinase GluC. Fragment ion annotation was performed using the interactive peptide spectral annotator^46^ with residues indicated in lower case p and d entered as hydroxylated residues (M+16), and in lower case d n as two residues that were hydroxylated and nitrile containing (+16 and −19 to give a net change of −3 Da).

Marfey’s analysis was used to determine the position and stereochemistry of the oxidation.^44,45^ After hydrolysis of the PesHI-modified peptide, the hydroxylated aspartate was reacted with 1-fluoro-2-4-dinitrophenyl-5-L-alanine amide (L-FDAA) and analyzed by liquid chromatography coupled to mass spectrometry detection (LC-MS). The product coeluted with an L-*threo*-3-hydroxyaspartate standard that was derivatized in the same manner (Figure 3a and S7). Collectively, these observations demonstrate that PesHI facilitate stereoselective hydroxylation of Asp21.

**Figure 3.**
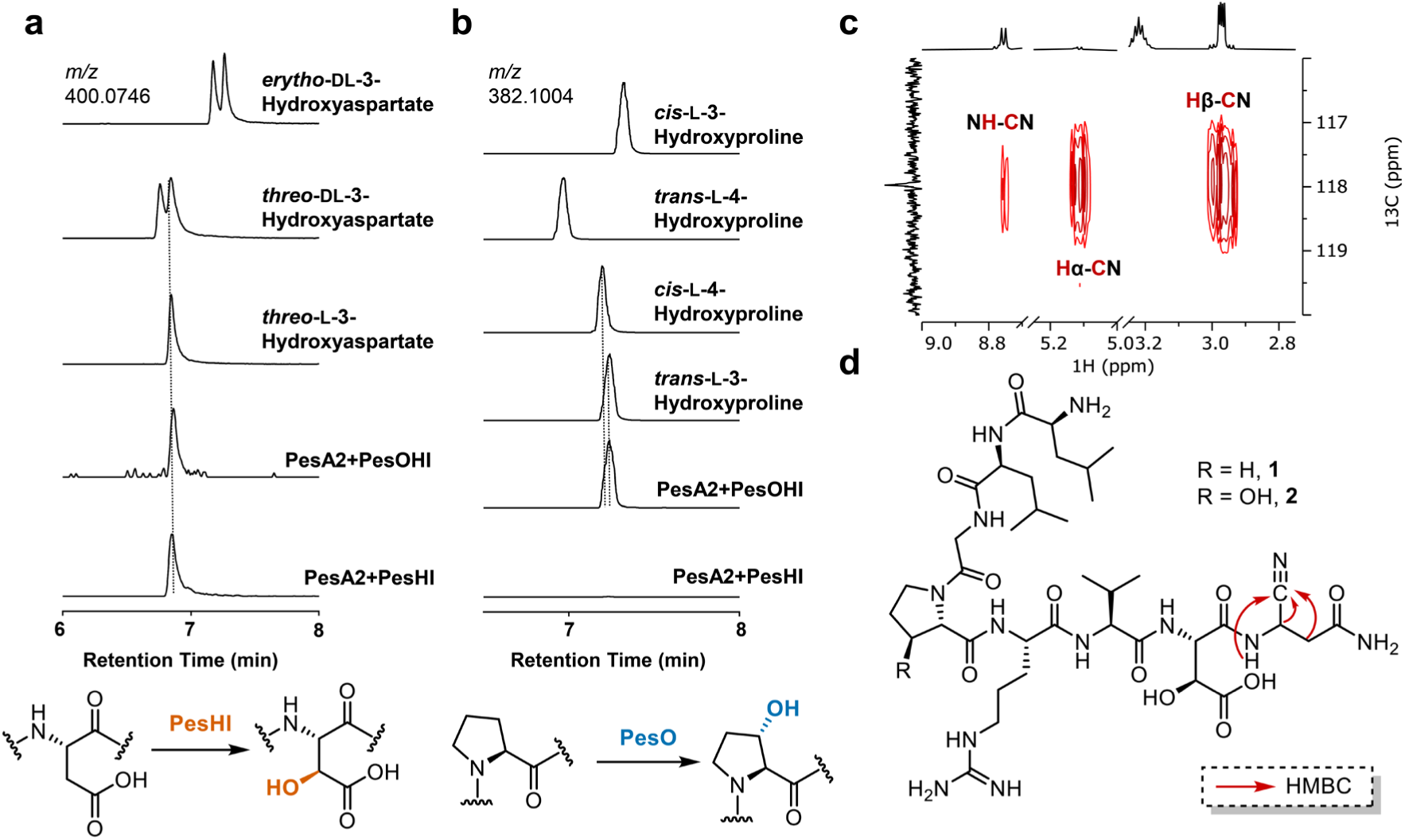
Structural determination of PesA2 co-expressed with PesOHIC. **a)** LC-MS analysis of hydroxyaspartate authentic standards or hydrolyzed PesA2-PesHI and PesA2-PesOHI peptides derivatized with L-FDAA; for coinjections, see Figure S7. **b)** LC-MS analysis of L-FDAA derivatized hydroxyproline from authentic standards or hydrolyzed PesA2-PesHI and PesA2-PesOHI peptides; **c)** ^1^H−^13^C HMBC spectrum of digested PesA2-PesHIC highlighting the C-terminal residue, and **d)** proposed structure of digested PesA2-PesHIC peptide **1** and PesA2-PesOHIC peptide **2** based on IR, NMR, MS/MS and Marfey’s analysis. Key HMBC correlations from **c)** are labeled.

Next, we included the αKG-HExxH protein PesO in the co-expression system in addition to PesHI, for which we will use the designation PesOHI. LC-HRMS analysis of the endoproteinase GluC-digested product peptide PesA2-PesOHI revealed the dominant production of a M+32 Da species (Figure 2). This additional +16 Da mass shift compared to the PesHI modification was assigned to Pro18 by HR-MS/MS analysis (Figure S6). We again performed Marfey’s analysis on the PesOHI-modified peptide to elucidate the identity of the hydroxyproline residue (Figure 3b and S7). Coelution with L-3*S*-hydroxyproline derivatized with Marfey’s reagent indicated that PesO stereoselectively hydroxylates carbon 3 of Pro18.

Incorporation of the AS-like enzyme PesC into the co-expression system resulted in a new product with a −19 Da mass shift relative to the PesOHI modified peptide. HR-MS/MS analysis demonstrated that a −3 Da mass shift had occurred to the C-terminal Asp21-Asn22 sequence (Figure 2 and S6). Asparagine synthetases usually catalyze the amidation of aspartate to yield asparagine,^47^ while in natural product biosynthesis, members of this enzyme family also catalyze lactam formation.^48–50^ Therefore, we envisioned that an amidation reaction of a carboxyl group (−1 Da) and a dehydration event (−18 Da) would account for the observed overall mass shift of −19 Da. When coupled to the PesHI catalyzed hydroxylation of Asp21 (+16 Da), amidation and dehydration would explain the net −3 Da shift of the Asp21-Asn22 motif. Within this sequence, the dehydration could originate in a dehydroAsp, or in the formation of ester, imide, or nitrile groups (Figure S8). The tandem MS/MS data were inconclusive in distinguishing these possibilities, and we therefore turned to analysis by nuclear magnetic resonance (NMR) spectroscopy described in the next section.

We also integrated the hypothetical protein PesX into the co-expression system. Co-expression of PesX in any combination with the other enzymes did not lead to the emergence of any new species derived from PesA2. These results imply that PesX was either non-functional in the *E*. *coli* heterologous host or the protein serves a non-catalytic role in the *pes* BGC. As described below, a potential function of PesX is suggested from *in vitro* experiments.

As noted above, co-expression experiments of PesA1 with Pes enzymes yielded similar modification patterns as PesA2 (Figure S9), indicating that the less conserved precursor N-termini as well as the residue at position 20 (Thr/Val) have minimal impact on the function of the modifying enzymes within the *pes* BGC.

### Structural Elucidation of Modified PesA2

The data described thus far suggest that PesC catalyzes consecutive amidation/dehydration reactions on the precursor peptide that is hydroxylated by PesHI on Asp21, yet the precise form of dehydration remained unclear. For NMR structural determination, we first attempted to process the modified PesA2 peptides with the native protease PesP1/P2. While the expression of this heterodimer in *E. coli* yielded soluble protein (Figure S10), prolonged incubation of the peptides and PesP2/P1 led to a range of proteolytic products, with a 25-mer peptide being the major product (Figure S11). The poor *in vitro* activities of PesP2/P1 limited their application in this study, and therefore we mutated Gln14 to Lys in PesA2 with the aim of generating a short peptide upon digestion with endoproteinase LysC (for residue numbering, see Figure 1a). We first explored the impact of this mutation on the PTM process, as well as the minimal enzyme requirement for complete C-terminal modification. HR-MS/MS analysis of this Q14K mutant after co-expression with PesHIC verified that the product still contained the −3 Da change in the Asp21-Asn22 sequence (Figure S12), indicating that Gln14 is not required for PesHIC modification. Therefore, we conducted large scale preparation of PesHIC-modified PesA2-Q14K in *E*. *coli*. After IMAC purification and LysC digestion, the resulting octapeptide (peptide **1**) was purified by high-performance liquid chromatography (HPLC) and used for NMR spectroscopic analysis.

A combination of 1D and 2D NMR experiments was conducted to elucidate the structure. Based on interactions observed in ^1^H-^1^H TOCSY, ^1^H-^1^H NOESY, ^1^H-^13^C HSQC and ^1^H-^13^C HMBC spectra (Figures S13-S15), all proton and carbon signals from each of the eight residues were assigned (Table S1). Importantly, all side chain protons and carbons of the Asp21-Asn22 sequence displayed chemical shift deviation compared to unmodified Asp/Asn residues. For Asp21, the β-carbon shifted downfield to 71.2 ppm and had only one proton attached to it, affirming that PesHI catalyzed β-hydroxylation of Asp21. Thus, the NMR data ruled out dehydration involving the hydroxyl group introduced by PesHI for the reaction catalyzed by PesC. The α-proton of Asn22 shifted downfield to 5.13 ppm, indicating a more electron-withdrawing environment than in a typical Asp. Together with the overall mass change, these findings are consistent with conversion of the C-terminal carboxylate into a nitrile group. Consistent with this hypothesis, the ^13^C NMR spectrum showed only 10 rather than the expected 11 signals in the range of 150-220 ppm (10 amide/carboxylate carbonyl carbons and one guanidine carbon), and instead a characteristic nitrile carbon signal at 118.0 ppm was observed. In the ^1^H-^13^C HMBC spectrum, this carbon signal showed correlations with the α and β protons and the backbone amide proton of former Asn22 (Figure 3c). Similar ^1^H and ^13^C chemical shifts have been reported on synthetic peptides that harbor C-terminal nitriles formed from _Asn.51,52_

We also isolated and characterized the peptide PesA2-Q14K after PesOHIC modification and LysC digestion by NMR spectroscopy (peptide **2**; Figures S16-S18). Compared with the PesA2-PesHIC peptide **1**, the signals characteristic of a β-hydroxyaspartate as well as the C-terminal nitrile group were retained, and additional chemical shift deviations from unmodified residues were mainly observed for protons and carbons associated with the Pro residue (Table S2). In the ^1^H-^1^H COSY and ^1^H-^13^C HSQC spectra, the observed correlations clearly assigned the hydroxy group to the β position (C3), consistent with the conclusions from Marfey’s analysis. This modification resulted in the conversion of a CH_2_ group to a CH group, accompanied by a downfield shift of the β-proton to 4.43 ppm and the corresponding β-carbon to 73.2 ppm. We also examined peptide **2** with Infrared (IR) spectroscopy, which revealed a characteristic triple bond stretch at 2150 cm^-1^. This signal was not present in the LysC digested, PesOHI-modified PesA2-Q14K peptide (Figure S19). The frequency deviates from a typical C-N triple bond stretching frequency (∼2200-2270 cm^-1^), perhaps because of interactions of the nitrile moiety with its environment.^13,14,53^ A ketenimine or isonitrile, which can display IR frequencies in the observed region,^54^ is inconsistent with the NMR data. Collectively, the IR, NMR and MS experiments, ^15^N labeling (see below), as well as precedent of nitrile formation by other ATP-dependent enzymes^17^ are consistent with PesC-catalyzed conversion of the peptide C-terminal carboxylate to a nitrile group.

### Characterization of the Reaction Catalyzed by the Nitrile Synthetase PesC

We next examined the PesC reaction *in vitro*. For typical AS-like enzymes, the N-terminus is buried inside the enzyme.^47^ Therefore, we constructed a PesC-encoding plasmid with a C-terminal His_6_-tag for expression in *E. coli*. In the canonical AS catalytic cycle, L-aspartate is converted to L-asparagine via ATP-dependent adenylation of the side chain carboxylate, followed by amidation using L-glutamine as the ammonia donor.^47^ Thus, IMAC-purified PesC was incubated with Mg^2+^, ATP, L-glutamine and the PesA2-PesOHI peptide for 6 hours. LC-MS analysis of the GluC-digested product demonstrated the formation of the C-terminal nitrile (Figure 4a). When ATP was omitted from the reaction, nitrilation activity was abolished. When the assay was performed for 1 hour, a product with a mass shift of −1 Da was observed by high resolution electron spray ionization mass spectrometry (HR-ESI MS) (Figure 4a) consistent with the formation of an intermediate C-terminal amide. To confirm the origin of the nitrogen atom in both products, the assay was also performed with L-glutamine-(*amide*-^15^N). The resulting C-terminal amide and nitrile containing peptides were both detected by LC-HRMS with a +1 Da mass shift (Figure 4b), indicating that in the PesC reaction, the inserted nitrogen is indeed derived from the side chain amide nitrogen of glutamine.

**Figure 4.**
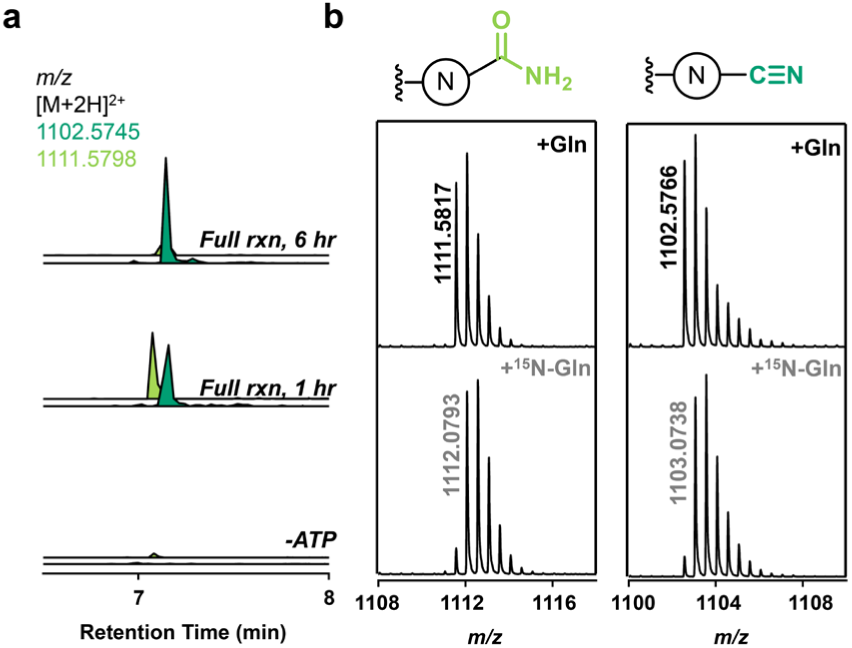
**a)** LC-MS analysis of the *in vitro* PesC reaction with PesA2 that had been co-expressed with PesOHI and treated with GluC (assay conditions: 100 μM PesA2-PesOHI peptide, 25 μM PesC, 5 mM ATP, 10 mM L-glutamine, 20 mM Mg^2+^); **b)** HR-ESI MS analysis of the resulting amide or nitrile species when L-glutamine or L-glutamine-(*amide*-^15^N) was used in the assay (calculated mass for unlabeled amide product [M+2H]^2+^: 1111.5798, unlabeled nitrile product [M+2H]^2+^: 1102.5745).

To investigate the leader peptide dependence of PesC, we purified PesOHI-modified, endoproteinase LysC digested PesA2-Q14K as well as PesOHI-modified, endoproteinase AspN digested PesA2. Reaction of the C-terminal 14-mer (AspN product) with PesC under the standard reaction conditions resulted in nitrile production (Figure S20). The C-terminal 8-mer (LysC product) was converted by PesC to the corresponding amide with partial conversion to a nitrile (Figure S20). These results suggest that PesC can still function in the absence of a leader peptide, but the attenuation in PesC efficiency suggests that the enzyme likely acts prior to proteolysis in the biosynthetic pathway.

Next, we explored the substrate requirements of PesC. The importance of prior PesHI/PesO modification for PesC activity was first examined. When unmodified precursor PesA2, PesA2-PesHI, and PesA2-PesOHI were incubated individually with PesC, the production of the corresponding nitrile products was only observed with PesHI-and PesOHI-modified PesA2 (Figure 4a, Figure S21). For unmodified PesA2, only trace amounts of the amide species were formed. Thus, the hydroxylation of the adjacent Asp residue on the substrate is essential for efficient amidation and for dehydration of the intermediate amide to occur, while hydroxylation by PesO is not required. This conclusion is consistent with the gene composition of orthologous BGCs, because *pesHI* homologs always co-occur with *pesC* homologs whereas *pesO* is only partially conserved (e.g. Figure S4). We also tested the tolerance of PesC toward alternative nitrogen donors. Similar to canonical asparagine synthetases,^55^ PesC also facilitated nitrile installation utilizing ammonia instead of L-glutamine as the nitrogen source (Figure S22b).

With the aim of potentially uncovering more AS-like enzymes that are capable of installing nitrile groups on ribosomal peptides, we performed a bioinformatic analysis on PesC. A BLASTp search with the PesC sequence as query only recovered high-identity homologs of PesC located in orthologous clusters of the *pes* BGC. Other less highly conserved hits were not related to RiPP biosynthesis, e.g. AsnO in the biosynthesis of *N*-acetylglutaminylglutamine amide.^56^ We then constructed a sequence similarity network (SSN) of the asparagine synthase protein family (PF00733), and examined their co-occurrence with RiPP biosynthesis elements (Figure S23).^57,58^ This analysis identified ∼3,800 candidates, of which ∼3,500 were related to lasso peptide biosynthesis. Among the remaining enzymes, TsrC^59^ and ScdTA^60^ are known to catalyze amidation of the C-terminal carboxylate of their substrate peptides. The remaining ∼300 uncharacterized candidates suggest a potential underexplored functional space of AS-like enzymes in RiPP biosynthesis. These enzymes may catalyze amidation, nitrilation, lactam formation, or currently uncharacterized reactions.

### Structure Comparison of Nitrile Synthetases and AS-like Amidation Enzymes

Using structure predictions and prior X-ray structures, we next investigated potential PesC active site features that might be important for catalysis. We generated the ATP and Mg^2+^ bound structures of PesC as well as the peptide amidating enzymes TsrC, ScdTA and AsnO with AlphaFold 3. Through structural alignment with the AMP-bound structure of *E. coli* AS (PDB ID: 1CT9),^61^ the ATP binding residues of PesC were identified. These residues are highly conserved among the five asparagine synthetase-like enzymes. PesC has one residue, Tyr340, that is not found in the other proteins which contain Phe, Lys, Met or Leu at the corresponding position (Figures 5 and S24). To investigate whether this unique residue conferred nitrilation activity, the Tyr residue was replaced by Phe. *In vitro* assay of PesA2-OHI with the PesC Y340F mutant revealed that the replacement had minimal impact on the nitrilation activity of PesC (Figure 5a).

**Figure 5.**
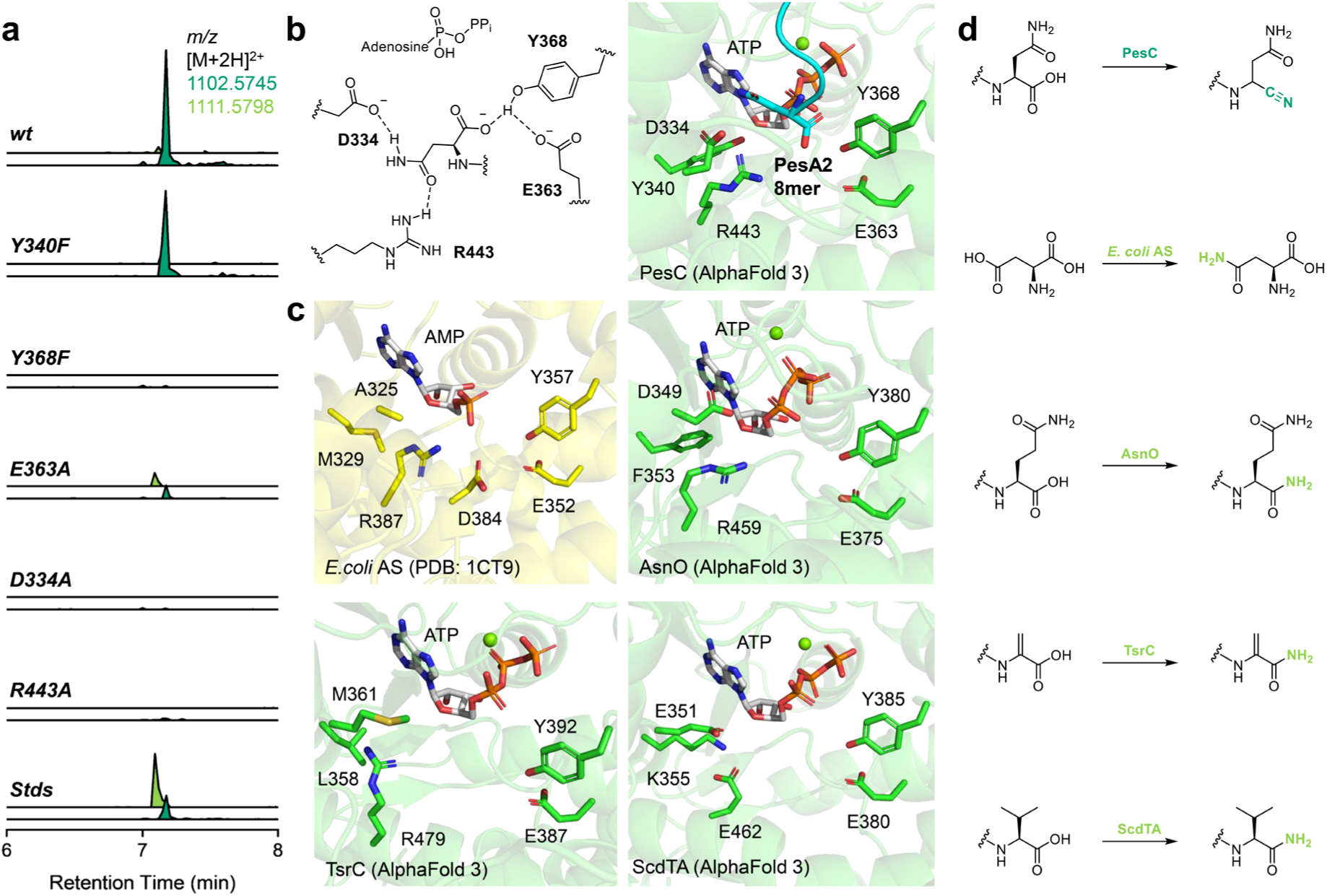
Structural and mutagenesis investigation of peptide amidation and nitrilation enzymes. **a)** LC-MS analysis of the *in vitro* reaction of wild type PesC and mutants with PesA2-PesOHI peptide after GluC digestion. The standard (Stds) trace derived from a 30 min PesC reaction with PesA2-PesOHI peptide when both amide (calculated mass [M+2H]^2+^: 1111.5798) and nitrile (calculated mass [M+2H]^2+^: 1102.5745) can be detected; **b)** The active site structure of the AlphaFold 3 model of PesC with ATP and the C-terminal 8mer of PesA2 and schematic representation of the proposed substrate recognition mode in the PesC active site. In the structural model, the C-terminal Asn of PesA2 is shown in cyan sticks, and active site residues that were examined by mutagenesis studies are shown in green sticks; **c)** Comparison of the active site structures of AMP-bound *E. coli* AS (PDB ID: 1CT9), AsnO-ATP (AlphaFold 3), TsrC-ATP (AlphaFold 3) and ScdTA-ATP (AlphaFold 3); **d)** Schematic representation of reactions catalyzed by PesC, *E. coli* AS, AsnO, ScdTA and TsrC. For the substrate peptides, only the C-terminal residue is shown.

Next, we examined the putative substrate binding pocket residues of PesC that are located near the ATP binding site (Figure 5). Glu363 and Tyr368 are conserved within all five enzymes, and are likely involved in recognition and activation of the carboxylate moiety of the substrates. In the AlphaFold 3 model of the unmodified C-terminal 8mer peptide of PesA2 bound to PesC and ATP, the C-terminal carboxylate of the substrate peptide is directed toward ATP through a hydrogen bonding network with Tyr368 and Glu363 (Figure 5b). Mutation of Tyr368 to Phe lead to complete loss of enzymatic activity, whereas the E363A mutant demonstrated reduced nitrilation activity as well as the accumulation of the amide intermediate (Figure 5a). Together, the Glu363-Tyr368 dyad likely plays important roles in substrate recognition and/or proton relay during PesC catalyzed nitrilation, and their conservation in all five enzymes suggest they are important for the common step of carboxylate adenylation. Asp334 and Arg443 are positioned across the substrate-binding pocket from the Glu363-Tyr368 dyad and may be involved in recognition of the amide side chain of the C-terminal asparagine of the peptide substrate. In *E. coli* AS, an Asp-Arg dyad is predicted to form hydrogen bonding interactions with the amino acid moiety of aspartate,^62^ and a similar arrangement is also observed in the active site of AsnO where the dipeptide substrate also possesses an amide side chain at the C-terminus (Figure 5b, c). Mutation of either Asp334 or Arg443 to alanine resulted in abolished activity, supporting a critical role of these residues in PesC substrate recognition (Figure 5a).

To gain further mechanistic insights into the nitrilation activity of PesC, we examined the ability of PesC to take an amidated peptide intermediate as substrate. We quenched the PesC *in vitro* reaction with PesA2-PesOHI after 30 min, when the amide was the major product (Figure S22a). After enzyme removal and buffer exchange, the resulting mixture was again incubated with PesC. Full conversion to nitrile was only observed when ATP was supplemented (Figure S22a), indicating that PesC catalyzed amide to nitrile conversion is also ATP-dependent. We also conducted PesC *in vitro* assay with the ATP analogs, *α*,*β*-methyleneadenosine 5’-triphosphate (*α*,*β*-AMPCPP) and *β,γ*-methyleneadenosine 5’-triphosphate (*β,γ*-AMPCPP). Conversion to the corresponding nitrile was only observed with *β,γ*-AMPCPP (Figure S22c), suggesting that the transformation of carboxylate to nitrile involves adenylation rather than phosphorylation of the substrate,^63^ consistent with other AS-like enzymes.

Based on these observations, a mechanism for the PesC catalyzed nitrilation can be proposed (Figure S25). The reaction is initiated by ATP-dependent adenylation of the C-terminal carboxylate, followed by the nucleophilic attack of glutamine-derived ammonia to form a C-terminal amide intermediate. The amide is then activated by a second equivalent of ATP to generate an AMP-imidate, from which AMP is eliminated to yield the C-terminal nitrile. This mechanism is similar to that previously proposed for ATP-dependent nitrile synthetases in other biosynthetic contexts such as the NRPS product auranthine and 7-cyano-7-deazaguanine (preQ_0_).^13,14,17^

### *In vitro* Characterization of MNIO Proteins

The co-expression experiments were unable to assign a role for the MNIO-like protein PesX. We therefore conducted *in vitro* characterization. Considering the prevalent partner dependence of MNIO proteins in the literature,^42^ we constructed PesHI and PesXI expression plasmids with a His_6_-tag on the N-terminus of either PesH or PesX. As anticipated, PesI was co-purified with both PesH and PesX (Figure 6b). Reaction of PesA2 with as-purified PesHI resulted in hydroxylation similar to the co-expression studies. When the assay was performed with PesH (no PesI) or with PesXI, no activity was observed by ESI-MS (Figure 6a). These findings are consistent with the absence of the iron-binding amino acids that are characteristic for MNIOs in the sequence of PesX (Figure S3), but do not provide any insights regarding the potential function of the protein.

**Figure 6.**
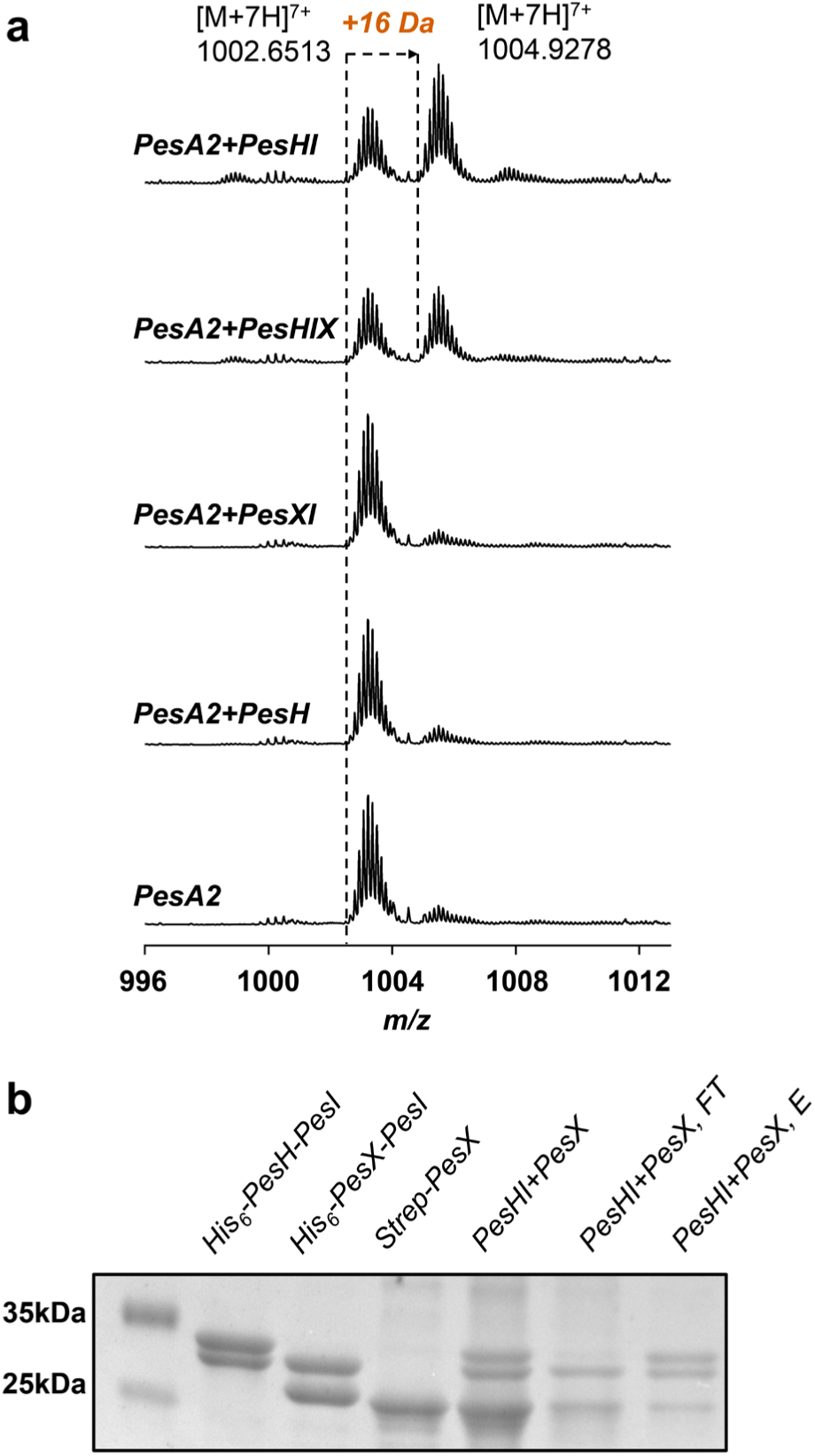
PesX inhibits the activity of PesH. **a)** ESI-MS analysis of the *in vitro* PesH/X/I reaction with PesA2 peptide (calculated mass for unmodified product [M+7H]^7+^: 1002.6476, hydroxylated product [M+7H]^7+^: 1104.9326); **b)** SDS-PAGE analysis of purified His_6_-PesH/PesI, His_6_-PesX/PesI, and Strep-PesX. Also shown are the flowthrough (FT) and elution (E) fractions of His_6_-PesH-PesI and Strep-PesX loaded on an IMAC column.

The ability of PesX to co-purify with partner protein PesI led us to speculate that PesX might serve to regulate PesH activity, and thereby nitrile formation. We generated structural models of the PesXI and PesHI heterodimers using AlphaFold 3, and alignment of these two structures revealed strong similarity (Figure S26). Similar to reported structures of the TglHI heterodimer,^64^ only the helical N-terminus of PesI interacts with PesH or PesX. The overlapping binding interface suggests that PesX can compete with PesH in binding to the partner protein PesI, which is critical for PesH activity as shown above. This proposal is supported by the observation that introduction of PesX in the PesHI activity assay resulted in diminished hydroxylation activity (Figure 5a). Moreover, addition of Strep-tagged PesX to a solution of the His_6_-PesH/PesI heterodimer, and subsequent addition to Ni-NTA resin resulted in the complex of PesX and PesI eluting from the column (Figure 5b). These observations indicate that PesX competes with PesH in binding with partner protein PesI, thus indirectly inhibiting its aspartate hydroxylation activity. Considering Asp21 hydroxylation is essential for PesC to perform nitrile formation, expression of PesX can slow down the production of the nitrile product. If correct, this regulatory mechanism suggests that the nitrile product may be toxic to the producing organism and that its production must be delicately regulated.

We performed bioinformatic analysis on PesX to gauge its distribution. A BLASTp search with an E value cutoff of 0.05 retrieved only PesX homologs from orthologous *pes*-like BGCs, suggesting that the sequence of PesX is highly specific to the nitrile formation pathway.

## Discussion

In genome mining campaigns for RiPP BGCs, a common strategy features a RiPP class-defining enzyme family as query.^25^ This approach has proven an efficient means to discover new and divergent compounds in existing RiPP families. Several ubiquitous families of RiPP post-translational modification enzymes are not currently identified as class-defining because they occur in many different RiPP classes. Two such enzyme families are the MNIO and αKG-HExxH enzymes.^31,42^ These proteins catalyze a variety of different transformations that at present cannot be predicted. In the current study, we investigated a representative BGC featuring members of both protein families that lead to the identification of the nitrilobacillins (Figure 7). Since the nitrile functional group is new to RiPPs and most likely functions as the pharmacophore of the mature products, we suggest the name nitrilotides for the wider group of products and the AS-like enzymes as the class-defining enzyme.

**Figure 7.**
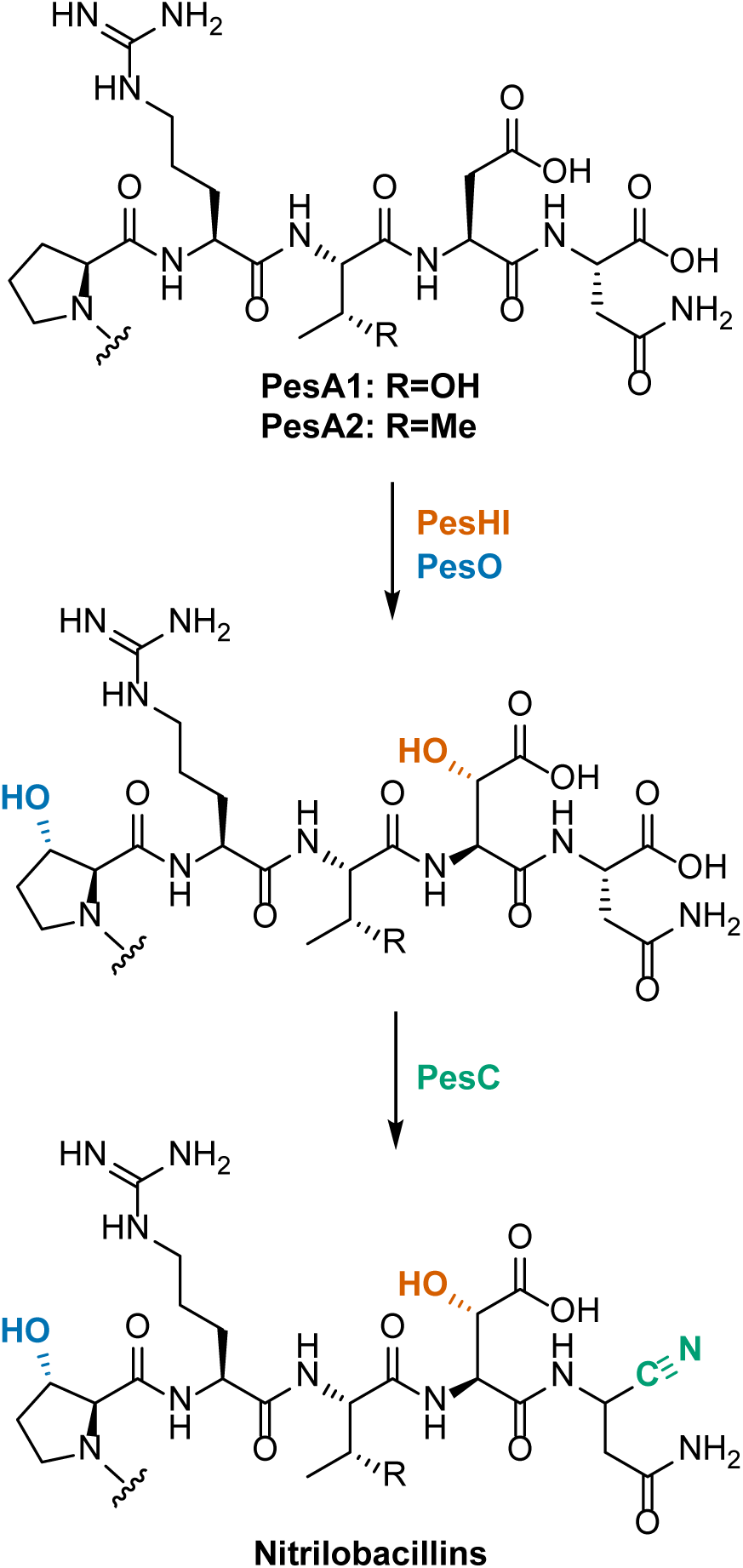
Proposed biosynthetic pathway of nitrilobacillins.

In medicinal chemistry, nitrile groups are commonly used as warheads including in in peptide-derived drug candidates. For instance, the nitriles effect covalent inhibition of cysteine/serine proteases in vildagliptin and nirmatrelvir (Figure S1b).^6,7^ Typically, peptidic protease inhibitors have structures at the P1-P3 position that resemble the native substrates. For instance, the pyrrolidone moiety at the P1 position of nirmatrelvir mimics the glutamine residue recognized by SARS-CoV-2 main protease.^18^ These observations suggest that the currently unknown physiological function of nitrilobacillins are likely inhibiting proteases that cleave at asparagine residues.

The AS-like enzyme PesC catalyzes the nitrilation at the C-terminal carboxylate in the PesA substrates expanding the previously reported small number of ATP-dependent nitrilation enzymes, ToyM/QueC^17^ and ArtA/NitB.^13,14^ During the revision of our manuscript, the characterization of another ATP-dependent nitrilation enzyme CalN, was reported that appears to have evolved from a kinase.^16^ Collectively, these enzymes introduce nitriles into nucleoside bases, polyketides and the side chains of amino acids. Other pathways to nitrile containing natural products involve oxidation of amino groups by heme proteins^9,10^ and flavin-containing monooxygenases (FMOs),^12^ as well as oxidative rearrangement processes of α-amino acids by non-heme iron dependent enzymes (Figure S1).^65^ Given the structures of their substrates, from a biocatalyst development perspective, PesC may present the most promising starting point for enzymatic installation of nitriles at the C-terminus of peptides. The reaction catalyzed by PesC has some differences compared to the precedented ATP-dependent systems. QueC/ToyM utilizes ammonia as the nitrogen source instead of Gln used by PesC, whereas ArtA/NitB and CalN catalyze only the amide-to-nitrile dehydration step. Crystallographic structures of QueC and ArtA have been solved,^13,66^ and alignment of the AlphaFold 3 predicted structures of PesC with the crystal structures of QueC or ArtA reveals that, even though chemically the transformation catalyzed by PesC and QueC are more similar, the overall fold of PesC is better aligned with ArtA (Figure S26). More specifically, the ATP binding domain of PesC exhibits structural similarity with that of ArtA. However, the substrate binding pocket of PesC is different from the active site of ArtA, and the key amide-interacting residues (Q146, S197, D199) identified through docking and mutational analysis of ArtA^13^ are not conserved in PesC. Instead, the active site of PesC adopts a much more open conformation in the model, likely to allow accommodation of large peptide substrates (Figure S26).

Enzymes from the AS protein family catalyze key transformations in the biosynthesis of bioactive natural products, such as β-lactamization in the biosynthesis of several β-lactam antibiotics.^48,62,67^ In RiPP biosynthesis, the lasso peptide cyclases belong to the AS enzyme family.^49,50^ In addition, several characterized asparagine synthetase-like enzymes such as TsrC^59^ and ScdTA^60^ amidate the C-termini of their substrate peptides. The nitrilation reactivity of PesC identified herein expands the diversity of chemical outcomes of asparagine synthetase-like enzymes in RiPP biosynthesis. Our current genome mining efforts identified hits that are likely involved in novel RiPP biosynthesis, but no homologs were predicted with high confidence to catalyze nitrile formation.

As in previous studies on ATP-dependent enzymes that catalyze nitrilation, the features that control enzymatic one-step amidation versus two-step nitrilation are not clear. Our *in vitro* PesC assays with amidated substrate as well as ATP analogs clearly suggested that both amidation and nitrilation steps involve ATP-dependent adenylation. Additionally, the nitrilation activity of PesC is not a processive reaction because the amide intermediate is observed and was converted to the nitrile product. The necessity of aspartate hydroxylation prior to nitrilation is unlikely to reflect a catalytic role during nitrilation because the absence of this modification also leads to nearly abolished amidation activity. The requirement for prior hydroxylation is therefore more likely involved in substrate recognition.

Site-directed mutagenesis experiments provided some insights into the reaction mechanism, which surprisingly is not well understood even for canonical asparagine synthetase B family members. Conservation of ATP-binding residues in PesC argues against a model in which an alternative ATP binding configuration enables nitrilation. The conservation of a Tyr-Asp dyad in all the enzymes in Figure 5 suggest these residues are involved in the only common step in catalysis, the amidation of the substrate carboxylate. Similar to reported examples of other tyrosine-carboxylate catalytic dyads,^67,68^ Tyr368 in PesC likely functions as the proton acceptor while Glu363 enhances the basicity of Tyr368 and assists proton relay during both amidation and nitrilation processes.

Overall, our combined mechanistic and structure prediction studies suggest that, compared with AS-like amidating enzymes, the unusual nitrilation activity of PesC does not derive from a single key residue, but more likely arises from a uniquely-tailored active site architecture that is perturbed to also favor nitrilation after initial amidation. Future structural studies may explain how the amide intermediate may be activated for a second round of adenylation.

αKG-HExxH proteins were recently identified to be iron and α-ketoglutarate dependent dioxygenases.^31^ Unlike canonical Fe/αKG dependent enzymes, proteins within this family utilize a conserved HExxH motif to bind iron, yielding a new active site architecture.^31,35^ The reactivity of PesO described in the current work is consistent with the reported β-hydroxylation activities of previous studies on members of the αKG-HExxH enzyme family,^31,35^ while revealing proline as a new site of modification since previous members hydroxylated His, Asp, Phe and Gln residues. Hydroxylations of proline at the β/γ carbons are common modifications in the context of single amino acid and protein side chains, and are usually catalyzed by canonical Fe/αKG enzymes.^69^ In RiPP biosynthesis, proline β,γ-dihydroxylation has been reported in the biosynthesis of microbisporicin and is catalyzed by the cytochrome P450 enzyme MibO.^70^

Multinuclear non-heme iron dependent oxidative enzymes (MNIOs) are an emerging class of metalloenzymes that contain two or three iron ions in their active site.^42^ Although this protein family comprises more than 14,000 members, only a handful have been functionally characterized. MNIOs are strongly associated with the tailoring of RiPPs. They catalyze a wide array of novel oxidative modifications to construct unusual scaffolds including oxazolones and thioamides,^36^ thiooxazoles,^38,39,71^ dihydroxyhomocitrulline,^72^ and *ortho*-tyrosines.^33^ Many of these are four-electron oxidations of their substrates.

PesHI is shown here to catalyze a comparatively simpler transformation, the stereoselective β-hydroxylation of aspartate to 3*S*-hydroxyAsp. This same two-electron oxidation reaction was also recently reported for an MNIO-nitroreductase fusion enzyme PflD where the MNIO domain catalyzes the transformation.^35^ In another related system, ApyHI catalyzes the oxidation of a C-terminal Asp residue to the corresponding alpha-keto acid with a β-hydroxylated aspartate as a proposed intermediate.^30^ Sequence alignment of PesH, ApyH and PflD revealed less than 30% identity for the MNIO domain. Notably, the conserved HxD motif that normally binds the third iron (Fe3) in MNIOs is missing for both PesH and PflD (Figure S3). Previous mutational studies on the MNIOs MbnB^73^ and TglH^64^ showed that mutating some Fe3 binding residues lead to decreased but not abolished enzymatic activity. Thus, the role of Fe3 in characterized MNIO enzymes remains elusive.

The function of the MNIO-like protein PesX is intriguing. No activity was detected *in vitro* or in *E. coli*, consistent with the absence of the metal binding residues in its sequence, which are conserved in all other MNIOs. Heterodimeric proteins composed of an inactive homolog with an active enzyme is not uncommon and is observed for instance with the Fe/αKG enzyme system CorB-CorD,^34^ and with several examples of metalloproteases.^74–77^ However, PesX appears to function in a different manner in that it competes with PesH for PesI, which is required for PesH activity. The identification of separate operons governing *pesH* and *pesXI* expression, whereas in other MNIO systems the HI proteins are often under control of a single operon (Figure S28), suggests that PesH activity is regulated in a more complicated manner. PesC catalyzed nitrilation only after hydroxylation of Asp21 by PesHI, suggests that this hydroxylation might serve as a gatekeeping event. The observed competition of PesX for binding PesI could therefore be relevant to tightly control nitrile formation within the native host, possibly to avoid toxicity. In this model, PesX and PesI would be expressed resulting in a non-functional heterodimer, which then would be converted to a fraction of active PesHI heterodimer upon initiation of PesH expression. Experimental test of this suppression strategy in species harboring *pes*-like BGCs requires investigations with native producing strains.

## Conclusion

In this work, we identified and characterized a novel RiPP BGC from *Peribacillus simplex.* The activity of the modifying enzymes was assigned through a combination of HR-ESI-MS/MS, NMR and Marfey’s analysis. MNIO and αKG-HExxH proteins were shown to perform stereoselective β-hydroxylation on aspartate and proline residues, respectively, and the Asn-synthetase-like enzyme PesC was demonstrated to catalyze two-step nitrile formation at the C-terminus of a ribosomally produced peptide. Mechanistic characterization of PesC identified key chemical and structural features required for nitrile installation. This study therefore expands the list of ribosomal peptide PTMs with nitrilation, and provides valuable insights into asparagine synthetase-like enzyme catalysis.

## ASSOCIATED CONTENT

### Supporting Information

Experimental procedures, Figures S1-S28 showing spectroscopic data, AlphaFold 3 modeling, and Tables S1-S3 listing NMR annotations and primers for PesC mutagenesis experiments.

## Funding

This manuscript is the result of funding in whole or in part by the National Institutes of Health (NIH; grant R37 GM058822). It is subject to the NIH Public Access Policy. Through acceptance of this federal funding, NIH has been given a right to make this manuscript publicly available in PubMed Central upon the Official Date of Publication, as defined by NIH. A Bruker UltrafleXtreme mass spectrometer used was purchased with support from the Roy J. Carver Charitable Trust (Grant No. 22-5622). W.A.v.d.D. is an Investigator of the Howard Hughes Medical Institute.

## Notes

The authors declare no competing financial interest.

## Acknowledgements

The authors thank Dr. Dinh T. Nguyen for helpful discussions, Matthew Albritton and Prof. Scott Denmark for help with IR spectroscopy and access to the PerkinElmer ATR-IR spectrometer, and Prof. Angad P. Mehta for access to an Agilent Synergy H1 plate reader.

## TOC graphic

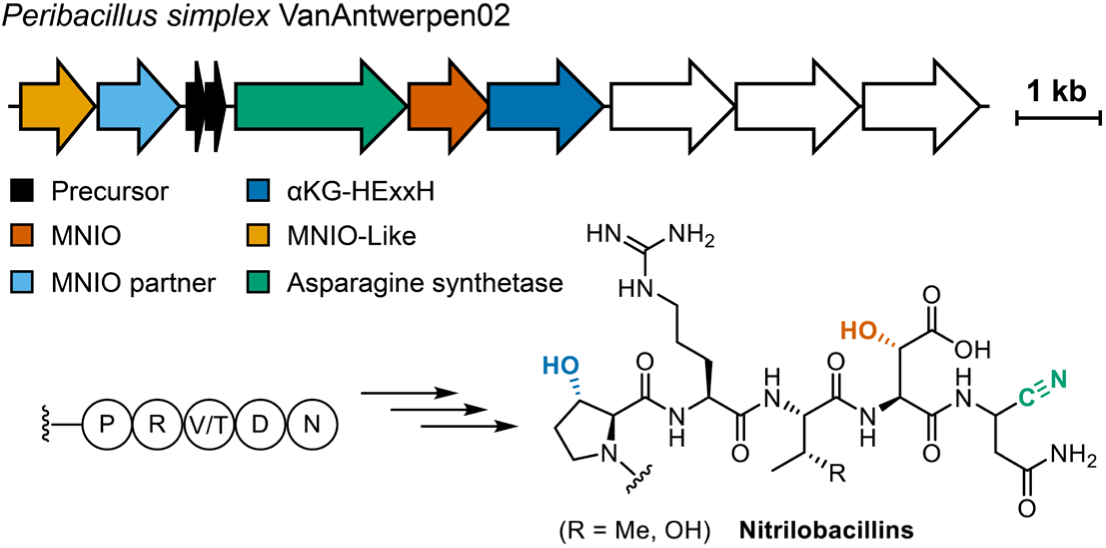

## Notes

### Competing Interest Statement

The authors have declared no competing interest.

### Summary of Updates

Additional experiments were performed to provide insights into the mechanism of catalysis of the nitrilation enzyme PesC resulting in new Fig. 5, and a new section describing these experiments. In addition, new figures were added to the Supp. Information (Figures S19, S20, S22, S24, Table S3) or were revised (Figure S3).

