## Supporting Information for "Discovery and Biosynthesis of Nitrilobacillins by Post-translational Introduction of C-Terminal Nitrile Groups"

**DNA constructs for expression and co-expression of *pes* BGC components in *E. coli.***

The DNA sequences encode the following genes (PesX: WP_155645489.1; PesI: WP_061142663.1; PesA1: WP_155645490.1; PesA2: WP_155645491.1; PesC: WP_061142664.1; PesH: WP_061142665.1; PesO: WP_061142666.1; PesP1: WP_061142668.1; PesP2: WP_061142669.1). They were codon-optimized for over-expression in *E. coli*, synthesized, and inserted into expression vectors pET-28a, pRSF-DUET and pACYC-DUET to obtain (co-)expression plasmids: pRSF-DUET::$His_6$-PesA1/A2(MCSI), pRSF-DUET::$His_6$-PesA1/A2(MCSI)::PesHI(MCSII), pRSF-DUET::$His_6$-PesA1/A2(MCSI)::PesOHI(MCSII), pACYC-DUET::PesC(MCSI), pET-28a::$His_6$-PesH/X-PesI, pET-28a::$His_6$-PesH, pET-28a::$^{C}His_6$-PesC, pET-28a::$His_6$-PesP2-PesP1 and pET-28a::Strep-PesX. The codon-optimized gene sequences are shown below.

*pesX*

ATGGAAGCAGGTGAAATTACAAAACGTATCGTGCACATTGAAGACTCGGACGTGG
CTCTTGACCAAGGGGTCCTGATTGGGGGGTTTCCAGTTGATTCCGTAATCTTCTCG
AACATTTTGTCTGATCGTTCTAAACGTGCTTCTCGCATGATTTATTCTCAATTCGGC
ATTAAGACAACGATTGCGTTACCGTATACGGACAAAGACTCTAAGTTGTTAGCAGA
TTATGTGTCGTTGCTCTCGGAACTTAACATTGACCAGGTGATTTTGAAGGTTGGTAC
CTGGGGTATCAATCATAGCTTCATGGACCCAGGGGGCTTCCAGCATCAGGATCAG
CAAAGTATTGAAGGCCTTCAGTCGTGGATCAGCGAGTTTAAGGAGTACCACGACAT
TCCGATCTATCTTCACCCAATTCGGCAACTGAGTCTTCCTCCGCAGTCCTTTTACAA
GAAAGTAATCACAGAACTTTCCGGGTTACATGTCCCGTTAAGTATTGATCTGGGCT
TGGTCTTATCCGATTGCAAAAATTGGGGCATGAAGTACAAAAAGGAGATCTTGGCG
CTTCTGCGCTCAAACCCGTCCATTTCCATGTTTTGGCTTTCGGCAGTACGCAAAGC
CGGGGACGCTTTCATCCCATGCGCTTGCGAATCTGTCGACGACCAAATTTGGGAA
ATGGTAGACTTGCTTATTGGTGAGGACGCTTCGTTGATTTTAATGACCGATCGTCT
GAGTAGTGAAACGCAGATTACCTCGGATGTTGAGAAGCTGATGAACGCAAAAGGT
GCAGTCCCGAAACATGAGTATTACGGGTGA

*pesI*

ATGAACACCACCGTAAAGCCGGAACTTGCGCTTCAAGACCTCCTCGTCAAAATGTC
CATCGATCCGAAGTGGTCTCCTGATATCAAGGATGGCAAGTTCCAACGGACTGTG
AAGAAAATGAACAAGGAAGAGCTTTTGCAACTGAACGTGAGTGCATTCCACTCGGA
GCGGAAAGGTCGGTGGCTGCAAATTCTTGCGCATCTGATCCCGATGTTCAGCATG
GGGATCCCTCTTCTTAAAGATGAGCTTAAATGCTCTTCAGACAATGAGCTCATGGA
AACCGATGTCTTCAACGAGTTCTTTAGTAGCCCTTTCCTCTGGAATATCGAGTTTAG

CATTCCGCACCCTCTGGGTTATGGTAAAGGCATCGAAAAAGCAACTGCTTTTATTT
ACTTCCTTCGCTCGTACAAGGAAATGTTGCCGCTCAAGCTCGTCGAGCTTTTCGAA
TTAGAATATGCAATCTTCAAGGCATCATATAAACTGCCATCAAAGAACACGTCGTTC
CTTGTAGGCAAACTCTACAAAAATACGTCGGCAACAATCATGAAGACTAAATACGA
TGTGCTCAGTGTTATTGAGGGTTGCGCTGACTTTCACGAATTGCTTCCGTGTGAAT
CGTTCTACATTATTTCTCCATTGAATAAAACGTACCGCATTAATTCGGATTATTACCG
TCTTCTGGCGTCTTTTGAGGAAGGCCATGAGACTGACAACTTGACTGACGGTGAG
CGTAACATCATCACGTCGGCTCTTAAGCTTCGTATTTTGTGTGCAAATCCAGTGGAT
ATTGAAGAACAGCTGATCGCACACAAGCGGTAA

*pesA1*

ATGGATAAGTCGATTAAGAATCAGGACTTGCAAACAGATAAAAAGGAACTCTCGCT
TAACGAGCTGCAAGAACTGGTTGCCCCAGGCTTTCTGGAGGCCAACCCGAACGGC
GGGCTGACTGATGCCGTCTTGGGCCAGTTATTAGGTCCTCGGACCGATAATTAA

*pesA2*

ATGAAGAACACGAAGAAGAACCCTGAACTTAACAGCGACAACAAAGAGTTGTCATT
AAACGAGCTCCAGGAGTTAGTGGCACCGGGTTTCCTCGAAGCAAACCCTAATGGC
GGGCTCACCGATGCGGTTCTGGGGCAACTGCTGGGGCCTCGCGTTGATAACTAA

*pesC*

ATGTGCGGGATATCTGGTTTTTTTGGACGCGAGGATAACAAGCCAATATACGTTAA
GACGTGTTTGAATGAAATGGAGCACCGTGGGCCGGACGATTGTAGTATCGTGGAG
TTGGATGAGATAGTGATGGGGACAGTGCGGTTATCCATACGTGACTCGTCAGAGA
AAGCTAACCAACCATTTGTAAGTAGTTCGGGGCGCTGGATTATCACGTTCAACGGC
GAATTGAATAATTTCCAACAGCTTCGCCACGAGTTGAATCGGTGCTGGGTCACCGA
ATGCGATACGGAGGTATTGGCGGAGTTATTTGAAACGTATGGGAAGGAAGGGGTA
CGCAAGCTGAAAGGAATGTACGCGATTGCGGCTTTCGACACGTACACAGACAAGC
TGTTTTTAGCCCGTGACTTCCCAGGTATTAAACCCTTATATTGGACTAAAACAGAGA
TTGGGGCCTATATTTACGCAAGCGAGTTTCCAGCATTGTTGAACACCCTTAAAGCA
CTTGACGAGGATTGTAAGATGACCGAAGCCCAGGTGTACGAATACCTTATGTACCG
TGATACTCTGCTTGAGCCTAACACGCTTATAAACTCTATAAAAAAGGTCTCACCAGG
AGAAATATTGGAGTTGAGTTTGGACAAGGAGGCCTTAAGTGATAAGATGAGTGGAG
TTCACGAGAAGAACGAGCCTCTTGATCTGGAAAGATTGTTTACCAAAGCGATCACA
TCGGTGATAGAATTGAAGCAGAAAATGGGTATGTTTCTGTCTGGTGGCGTCGATTC
AAGCACCGTGCTGTCAGAGATGGTATCGCAGGGTGCAGAGGTGGAGGCGTTTAC
CTTGAGATACGAAAAGACTGGAGAATTCTCTTACTCCGAGGTGCCTTTTTCAAAATA

CGTTTGTGAATTTCTGAATGTCCCTTTGCATGAGGTTATCCTTACCGAGAAAGATTT
CATCGAGCTTCTGCCCCAGGCACTGGCGAACCAAGATGGGCTTAGCATGGACCCA
ACAATAGTGGCATATTACGCCTTGGGTGAAGCAGCCGAGAAGAGAGGGCTTAAAG
TGGTGATCACCGGGACGGGGTCCGACGAGATATTCGGCAGTTACGATTGGCTTCA
TTCAGATTCGATAGAGCGTTTCGATCATTGGATGAAGTCTGCAACGACTAAATCTCT
GTTGCGCATGGATGAGCAAGTATGGAAGAATATATCCGACTCTTCATTTCTTGACC
GGAAGCATAAGATAGAGACGTTCATGTCCGTCGGCGGGAATTTGGGGGACGCAAT
TAGATTGTTTGGGTTCTATCACTTAGAAGCAGATGCGTTGCCTCGTGTGGACCTGG
GGACAATGAATAGCTCCGTTGAGTGTCGGGTCCCCCTGCTTGATCAAGACTTCGTC
CAAGCCGCTCTGTCTATGTCCCCAGATAAGGAAAAGTCCTTATTCCGGAGTTTGTC
AAAGAATAGAATACCCATTGAGATTCAACAACGTCCCAAGTGCGGGTTCCCACATC
CTGTTTTCATTTGGATGAAACATGGTCAACTTGGCAGAGTAGCATATGACTTTTTGA
AGATATCGCAACTTAAGGACGTAATAAACTTTGAGGAACTGGACAAATTCTTGAAC
GATGGAAGTAGTCTGCTGGGACGGAATCCTTCGGCCAGACTTTATGATGTGGGCA
GTAGCTTTTGGTATATCTACTCGTTCGCCCTGTGGTGCGAAATCCATAAAGTTCGTT
TACCCAATCTGAAAAAGTGCAAGGAAGTTTATTATGAGAATTGA

*pesH*

ATGAAAACCGAAACTCAACTCGCGAACGAGGTTATTAGCGGGAAACGGTTGGGTA
TTGGTGTTGCCTGGATCGACAGCCCAGAATACCTTGATTTTATTAACAAAGAGCTT
GACCCGAGTTTCCATTACATCGAGGTACACTTACCTAGCAATATTAACAATGTTCCT
TCTAATTTCGAGAAAATTGCCCACTGTTCAACATTACCACTCGCTGACTTACACGAA
CCGACTCGCAAAATCATTAAAGACGTCTTCCATCAGGCGAAAGAAATCGAGGCGC
ATTGGATTGGGGAGCACCTTTCAATTCTCGGCACCACTGACGGCCTTCAGTTTGGT
TATATCTTTGCGCCAATCAAAGACGAAGACTTGAAAGACCGCGTTATCCGGCGGAT
TAAGCGCTATCAAGAAACGTACGGTATTCCGTTTGTTATTGAATTAGGTCCGCGCT
ATCACGATTGGGGCGGTGTTGACGTCTTGGAAGATTACACGCTTCTTAAGGAAATT
AGCATTGAGGCGAATATCCCAATTATTCTGGACATTCCACATACATGGGCCACCGC
TAAAGCATTCAATAAGAGTTTTGAAGAACTTTTATCGTTCCTGAAAGGCGCGCACAT
CGCGGAGATCCACATCGGGGCATCAGGTGTAAGCAAGAAGGGGCAAATTTCTTTC
GAGCAATCTTGGAACCAGCTGGTAATCTGCCTGGAAATGTTCCCTGAGGTGCGCG
GTATCACTGTCGAATTAGGGAAAAATACCAAGACGGAAGACTACTGTAATACAATC
CGCGAGGTCAGTAAACGGTTCCAATCGTCCAAATGGCGTATCAATGACGAGAACA
ATTGA

*pesO*

ATGAAAACAATCGACTTTAAGAACTTACTCACACCAGCTCCGTCGAATATCGGGCG
TGACGTTCATAAGCAGCTGCTTAGTCTCCGTAATATCAAAGCGCTTGAGCTCATTTT

GCAGTCGAACAAGGAGGAGAATCTTAACTGGCTTGGTAAAGCCATTGATTTGCTTA
AAACGTGTGATGCAGCCGAACGCGACCACGTGCTGAACAAGCCGTATGTCTATCT
GTGGATCGAAAAATATAAGAATTTCACATCGATGGGTGAAACTGAGAAGCAAACTT
GGATTTGGAAGGGTTTCGTGCACATCTTCTTACAGTTAAACAAACGGGTCAATCCT
GTGTTGGAATCCTTACGTCTTCGCTTTACCGTTAAGCCTAAAGACATTTTCTTCCTG
AAAGATATGACTGAGAATCAGGCCGCTAAGGAGATTCTTGTTATCAGTGAAAATAA
CCACATTACTTTCGAGGTCGAGCACGTCCTGTACAGTTTTGATGTCCATACATTCCA
CAAAGCCTTTACACTGCCGTTCATCACGAACGAAGTATTTTGGTTTGAGGAGTACG
TTAACATTGCCAGTTCTGCCGAATTACGTATGAATATGGAGATCACAGAAGTTGTCA
GCGAGGACATCAACAATTCGTTTGAAGAAGCCCTTCAGCTCATCAAAATTAGTTGG
CCTGAGATGTTTGCAGAAATCGATGGCAATATTGAAAACATCGTTTTCTTTAACCCA
CCAGGCTTGCCGTTTTTCTCGGACTTCCGTGTCCACGGCACTATTTTTGCTTGTCA
TGTGGCACGGCCGGTAGTGAAGATTGCCGAATGGCTTATTCACGAGGCGTCCCAC
AATCGCTTAAATACAATTATGTGTGCACGGCCTCACATTTTTAATGACATGAAACCA
ATCTATACCAGTCCATGGCGTGACGAGTTGCGCCCATTATACGGGATTTATCACGG
TTGTTTCGTCCATTGTCGGGTGCTCCACTTCTACCGGCTCTTAAAGAAGAAGCGTG
CTACATACAAAGGTGTGTCGATGGAAAGTGAAATCGAACGGATCTCGGAAGAGTT
GCAATTAGGTTTGAGCACTATTGAGGAGCATGCGTCCCTCTCGCCTGACGGGGAA
GCCCTGGTCCATGGGATGCAAAACTTGGTTATCGCGGACAGCAAGTGA

*pesP1*

ATGAGAAATGACTTCGTGACTCGGGAGCAAGGAAACTTTTCGTTTCACCTTTTACC
CTCGGAAAAATACGTTCAGACAACTATAGCGGTTAGAATTCACTTGGATCTTGACG
AAAAGATGACAACCGGCGCCGCACTGTTGCCATACATTCTGTTACACGGCAGTGA
ACGTTTCCGGAATGACTTCATAATCCAAAGTAAGCTTGATGAGGAATATGGAGCAA
AGCTGGGTGTTAGCATAGACAAAAAAGGCGACAAGCAAGTCTTATGCTTATCCTTG
AAGTTTTACAATCAAAAACACAAAATGATTAAACCGGAAATACTTACTGTGTTACTG
GACTTGCTTACTCAACCATTAATTACGAAAAAAAGTGTGGAACTGGAGAAGAAGTT
GCATTCCCGGCGGATTCAAAACGAAATATTCAATGCTTACCACTATAGCTACAAAAA
GAGCTTTTTATATCTTCAAGGTAACCAACAAAAGTTGTACTTGGACCAGGTGGGGT
CTATCGGAGACATCGAGACATGGAACGAATCGAAATTACGCAACCTGCACCAGAA
GATCCTGCAAACCGCCCCGATCCATGTTTATGTTATTGGAGATATTGACGAGGAGG
AAGTATTGGAGCAATTAAGTAGTTGCTTATGCGGGGGAGTAGAAGAATCCGAGCG
GAGCATGATACCGCCAACACCAGTGCAGTCTCCGGTTAAGACCTATCACTGCGAA
AAGTTCCAGAGTAAGCATGCTATCATTACTGCATGTAGTTCAACCGGTGGAATCAC
ACTTAAATCAGAGAAATATCCTGCCTTAGTTGTATTTAATCGGTTGTTGGGCGGATT
TCCACATTCTCGTCTGTTTATGAATGTCAGAAACCAATTACAAGCCGCTTACTCAAT
CATTTCTATTCTGGACGCCATTAATGGGTTACTTTTCATACAGACCGCTATTGATCC

GGCCTCGGAGGAACTTGTCTTACGCAAAATATTCGGGGAGATGAAGAGTATTAAAG
AAGGAGGATGCACGCAGGAGGAATTAGAAGCAACAATAAAAACATTGAAGAACTCA
TATAAGTTATCTCAGGACTCTCCCGTCTCGCTGATTGACTTCCATCAAAACGGCATC
ATATCTGGCAAGGTGCGCACCATGACCGAGATGATAGACTGCATTGACCGGATAG
AACTTCAGGACATCCAGGAAGTGGCTCAGGTGTTTAAAGGATACTCAATATTTACG
CTGGGTGAGGGGTGA

*pesP2*

ATGGTCATAGGTAGTCACATCGAGAAAAAAAATATAAACGGTTTGGACACGTTGCT
TGTAAATAAGAAAGGATTTTTGGAAAATTATATATTGTTAAGCGTTGGTTTTGGTGG
GATGGATAGTCACTTTCAGGACGGTTGGAACCGTAAGATCCCGTATGGTACTGCA
CATTTTCTTGAGCATCTGATTATTCAGAACGTCCAATTCGACGCGATATATCGCTTT
GGAGAAATAGGAGCATCACTTAATGCTTCAACTAATTTCGAGCGTACGAGCTTTAG
CATAAGTTGTACCGAACAGTTAGGATTGAACATTCAAATGATTCTGAAATTGGTCCA
GTCAATTTCCATTACAAAGGACATAGTAGAAAAGGAGAAGAAGATAATCAAGCAGG
AATTCATGCAATTGCACAGTAACCGGAAAAACTCCTGTTATACACACCTTCTGAGC
GATTTGTATGGCGGCGAGAGTGGTTTGGCGCAACCTATCATCGGAAACTCGGGGT
CCATTGAAAGTATGGATGCATGCATGCTTCAAAAATGCTTCCAAAAGTTTTACTCTA
TTGAAAATATGAAAATAATCATAATTGGCGAACTTAACCCTGAGGAGGTCTACGAC
GTGATAGAACGGGCGTTGGCGAATAGTCCCGTCAAAGAAAAAGAACGGTTCTCCC
CGTTCATACCGGTGAAGCGCGAGAGCTTGAAGATTGAACAGATAAATTCTACACAA
TCAGTGTTATTCTGGGGAAACGCGATCACCTACGGCAACGACGCAGGAAATCCCT
ATAATGATATTCGTAAGGAGTTTTCAATACGCGTGGGGTTGGAAATATTATTTGGAA
GAACTTCAGAGTTTCAGCGGAAGATACATGGCCAAGGCCTTATCGATGGGTCAATT
GATGTCAAATGCGAATTTTCTAACCGTTACTGTTTTTCACTTCTGAATACAATCACG
AATAAACCTTCTGACGTGATGGAGGAGATAGACAAGACCAAAAAAGATATACGCAA
GCGTTCAATTGGTGACGTAGAATTTCTGATTTCAAAAAGACGGATTATTGGCAGAAT
CATCGACGTGTATGATCATCCTGCTAAACTTGCAAACAGCATACTTCATTATAGCAT
AAGAGGCATGGATTTCTCCGACGTGCTTGACGTTATGAACTTAATCACGAAAAAGG
AAGTAATGGATGCTATGCATTCGCACCTGTTATAA

**Overexpression and purification of (modified) precursor peptides**

The above-mentioned precursor peptide and modifying enzyme (co-)expression plasmids were used to transform chemically competent *E. coli* BL21(DE3) cells. A single colony was selected from the Luria broth (LB) agar plate to inoculate LB containing 50 µg/mL kanamycin (and 20 µg/mL chloramphenicol depending on the plasmids), and the culture was grown overnight at 37 °C. The cultures were used to inoculate 2 L of LB medium with kanamycin (and chloramphenicol), and grown at 37 °C with shaking at 220 rpm until the

OD600 reached approximately 0.6. The cells were placed on ice for 30 min, supplemented with 50 mg iron(II) sulfate and 150 mg sodium citrate for MNIO-related co-expressions, and the expression was induced by the addition of IPTG to a final concentration of 0.4 mM. The induced cells were grown at 18 °C with shaking at 220 rpm for 14-18 h. Cells were harvested by centrifugation at 6,000 rpm for 20 min at 18 °C. The cell pellet was resuspended in 80 mL of lysis buffer (100 mM sodium phosphate, 300 mM NaCl, 10 mM imidazole, 6 M guanidine HCl, 10 mM Tris, pH 8) per 30 g of cell pellet, and stirred at 4 °C until the cell pellet completely resuspended in buffer. The cell membrane was disrupted by sonication, and the clarified cultures were centrifuged at 20,000 rpm for 40 min. The supernatant was loaded onto a 2.5 mL Ni-nitrilotriacetic acid (Ni-NTA) gravity column, which was pre-equilibrated with 5 column volumes (CV) of lysis buffer. 10 CV of wash buffer (100 mM sodium phosphate, 300 mM NaCl, 30 mM imidazole, 6 M guanidine HCl, 10 mM Tris, pH 8) was added to the column after the supernatant had drained completely. The desired precursor peptides were eluted by the addition of 5 CV of elution buffer (100 mM sodium phosphate, 300 mM NaCl, 300 mM imidazole, 4 M guanidine HCl, 10 mM Tris, pH 8). The eluents were concentrated to 2.5 mL with a 3 kDa centrifugal ultrafiltration tube. The concentrated eluent was loaded onto a Prepacked Desalting column (PD-10), which was pre-equilibrated with 5 CV of preservation buffer (100 mM Tris, 150 mM NaCl, 1 mM TCEP, pH 8). After the loaded eluent drained, 3.5 mL of preservation buffer was added to the PD-10 column, and the new eluent was collected. The collected eluent was centrifuged at 7,000 rpm for 5 min to remove precipitates, and the concentration of purified precursors was estimated by the nanodrop A205 protein quantitation method. Purified precursor peptide stock solutions were stored at -20 °C.

For large scale preparation of PesA2(Q14K)-PesHIC and PesA2(Q14K)-PesOHIC, the corresponding overnight cultures were used to inoculate 6 L of terrific broth (TB). After induction, the cells were incubated at 18 °C for 38-42 h. The desired peptides were then purified following a similar procedure as described above except that 10 mL Ni-NTA resin was used and all volumes were scaled accordingly.

**High-resolution tandem mass spectrometry of digested precursor peptides**

The peptides from Ni-NTA purifications were digested with endoproteinase GluC in a 1:20 mass ratio. After acidification and filtration, the digested peptides were injected onto an Agilent 1290 LC-MS QToF for ESI-HRMS and MS/MS analysis. LC separation was achieved at 50 °C using a Phenomenex Aeris 2.6 μm PEPTIDE XB-C18 LC column at a flow rate of 1 mL/min using 0.1% formic acid in $H_2O$ (solvent A) and 0.1% formic acid in acetonitrile (solvent B) with the following gradient program: 0–2 min 5% B, 2–11 min 5–80% B, 11–11.1 min 80–95% B, 11.1–15 min 95% B, 15–17 min 95–5% B. Mass spectra were collected in positive mode at 10 spectra/s and 100 ms/spectrum. Tandem-MS fragmentation was achieved at normalized collision energies of 20, 25 and 30. HR-MS/MS analysis was performed using the Interactive Peptide Spectral Annotator (IPSA) tool[1] and verified manually.

### Marfey's Analysis

The PesA1-PesHI, PesA1-PesOHI, PesA2-PesHI and PesA2-PesOHI peptides from Ni-NTA purifications were further purified using a Vydac protein & peptide C18 column (particle size: 5 µm; dimensions 250 x 4.6 mm) at a flow rate of 0.8 mL/min using 0.1% trifluoroacetic acid in $H_2O$ (solvent A) and 0.1% trifluoroacetic acid in acetonitrile (solvent B) with the following gradient program: 0–5 min 5% B, 5–35 min 5–100% B, 35–40 min 100% B, 40–43 min 100–5% B. Fractions containing desired peptides were collected and lyophilized. Around 1 mg of each purified peptide was hydrolyzed in 0.8 mL of 6 M DCl in $D_2O$ at 120 °C for 6 h. The resulting mixture was concentrated under reduced pressure, resuspended in 0.8 mL deionized water, and concentrated again. This mixture was then again resuspended in 0.8 mL deionized water and lyophilized. The hydrolysate was then dissolved in 100 µL water, and 50 µL of the solution was mixed with 100 µL L-FDAA solution (0.1% w/v, in acetonitrile) and 20 µL of 1 M $NaHCO_3$. The mixture was allowed to incubate at 43 °C for 1 h before 20 µL of 1 M HCl and 10 µL of $H_2O$ was added. The mixtures were subject to LC-MS analysis on a Phenomenex Aeris 2.6 µm PEPTIDE XB-C18 LC column at 50 °C at a flow rate of 1 mL/min using 0.1% formic acid in $H_2O$ (solvent A) and 0.1% formic acid in acetonitrile (solvent B) with the following gradient program: 0–2 min 5% B, 2–12 min 5–95% B, 12–17 min 95% B, 17–19 min 95–5% B. For co-injection experiments, the derivatized hydrolysate samples were mixed with standards in roughly 1:1 ratio, followed by LC-MS analysis. The DL-*erytho*-3-hydroxyaspartic acid standard was synthesized following a reported procedure,[2] and all other hydroxylated amino acid standards were purchased from Ambeed, Santa Cruz Biotech, Oakwood or Sigma.

### HPLC purification of protease digested modified precursor peptides

The PesA2(Q14K)-PesHIC and PesA2(Q14K)-PesOHIC peptides from Ni-NTA purifications were digested with endoproteinase LysC in a 50:1 mass ratio. After overnight incubation at 37 °C, the completeness of digestion was checked via MALDI-TOF MS. The fully digested mixture was then acidified, and filtered before purification by HPLC. HPLC purifications were conducted with an Agilent 1260 Infinity III system, and both peptides were purified with a Waters XBridge® Prep C18 column (particle size: 5 µm; dimensions 250 x 10 mm) at a flow rate of 2.5 mL/min using 0.1% formic acid in $H_2O$ (solvent A) and 0.1% formic acid in acetonitrile (solvent B) with the following gradient program: 0–5 min 5% B, 5–35 min 5–60% B, 35–38 min 60–100% B, 38–40 min 100% B, 40–43 min 5% B. Fractions containing desired peptides were collected and lyophilized. For PesA2(Q14K)-PesHIC core peptide, the lyophilization product was directly used for NMR analysis, while the resulting crude PesA2(Q14K)-PesOHIC core peptide was repurified with a Waters XBridge® C18 column (particle size: 5 µm; dimensions 250 x 4.6 mm) at a flow rate of 0.8 mL/min using 0.1% trifluoroacetic acid in $H_2O$ (solvent A) and 0.1% trifluoroacetic acid in acetonitrile (solvent B) with the following gradient program: 0–5 min 5% B, 5–10 min 5–20% B, 10–30 min 20–30% B, 30–33 min 30–5% B. The resulting pure PesA2(Q14K)-PesOHIC core peptide was lyophilized, and the buffer was exchanged back to solvents containing 0.1% formic acid with the above-mentioned semi-prep column and method. Then this peptide was lyophilized again and used for NMR analysis. The additional buffer exchange procedure mentioned herein helps prevent TFA carbon signals from interfering with the key nitrile carbon signal at ~120 ppm.

For the PesA2-PesOHI 8-mer and 14-mer peptides, full length PesA2 (Q14K)-PesOHI and PesA2-PesOHI peptides were digested with 50:1 mass ratio of LysC and AspN, respectively. After digestion, the desired peptides were purified with a Waters XBridge® C18 column (particle size: 5 µm; dimensions 250 x 4.6 mm) at a flow rate of 0.8 mL/min using 0.1% trifluoroacetic acid in $H_2O$ (solvent A) and 0.1% trifluoroacetic acid in acetonitrile (solvent B) with the following gradient program: 0–5 min 5% B, 5–35 min 5–70% B, 35–38 min 70–100% B, 38–40 min 100% B, 40–43 min 100–5% B.

**Overexpression and purification of proteins**

The above-mentioned His-tagged enzyme expression plasmids were used to transform chemically competent *E. coli* BL21(DE3) cells. The following day, a single colony on the plate was selected and used to inoculate Luria broth (LB) containing 50 µg/mL kanamycin and grown overnight at 37°C. The cultures were used to inoculate 3 L of LB medium with kanamycin (and chloramphenicol), and were grown at 37 °C with shaking at 220 rpm until the OD600 reached approximately 0.6. The cells were placed on ice for 30 min, and expression was induced by the addition of IPTG to a final concentration of 0.2 mM. The induced cells were grown at 18 °C with shaking at 220 rpm for 14-18 h. Cells were harvested by centrifugation at 6,000 rpm for 20 min at 18 °C. The cell pellet was resuspended in 80 mL of lysis buffer (50 mM Tris, 300 mM NaCl, 10 mM imidazole, 10% glycerol, pH 7.5) per 30 g of cell pellet with the addition of 1 mg/mL lysozyme and Pierce™ Protease Inhibitor Mini Tablet (EDTA free), and stirred at 4 °C until the cell pellet resuspended completely in buffer. The cells were disrupted by sonication, and the clarified cultures were centrifuged at 20,000 rpm for 40 min. The supernatant was loaded onto a 2.5 mL Ni-nitrilotriacetic acid (Ni-NTA) gravity column, which was pre-equilibrated with 5 column volumes (CV) of lysis buffer. After loading, 10 CV of wash buffer (50 mM Tris, 300 mM NaCl, 30 mM imidazole, 10% glycerol, pH 7.5) was added to the column. The desired proteins were eluted by the addition of 5 CV of elution buffer (50 mM Tris, 300 mM NaCl, 300 mM imidazole, 10% glycerol, pH 7.5). The eluents were concentrated to 2.5 mL with 10 kDa or 30 kDa centrifugal ultrafiltration tubes. The concentrated eluent was loaded onto a Prepacked Desalting column (PD-10), which was pre-equilibrated with 5 CV of preservation buffer (50 mM Tris, 300 mM NaCl, 10% glycerol, pH 7.5). After the loaded protein solution had drained, 3.5 mL of preservation buffer was added to the PD-10 column, and the new eluent was collected. The collected eluent was concentrated again with centrifugal ultrafiltration tubes, and the concentration of purified proteins was estimated by the nanodrop A280 protein quantitation method. Purified protein stock solutions were stored at -80 °C.

For the purification of Strep-PesX, the cell pellet was resuspended in lysis buffer containing 100 mM Tris, 150 mM NaCl, 1 mM EDTA at pH 8. After cell lysis and centrifugation, supernatant from 6 L LB culture was loaded onto 5 mL IBA Strep-tactin® 4Flow® high capacity resin. Then 5 CV of wash buffer 1 (100 mM Tris, 150 mM NaCl, 0.01% Triton X-100, 1 mM EDTA, pH 8) and 5 CV of wash buffer 2 (100 mM Tris, 150 mM NaCl, 1 mM EDTA, pH 8) was passed through the column before the desired protein was eluted with 3 CV of elution buffer (100 mM Tris, 150 mM NaCl, 2.5 mM desthiobiotin, 1 mM EDTA, pH 8). The subsequent procedures were same as above described except that the protein was exchanged into buffer containing 100 mM Tris, 150 mM NaCl at pH 8.

**NMR/IR data acquisition and analysis**

The PesA2(Q14K)-PesHIC core peptide sample was prepared by dissolving the purified peptide in 500 µL of 90% $H_2O$, 10% $D_2O$ with a final concentration of ~3.8 mM, while the PesA2(Q14K)-PesOHIC core peptide sample was dissolved in 500 µL of 90% $H_2O$, 10% $D_2O$, 0.1% $d_2$-formic acid with a final concentration of ~1.2 mM. NMR data were collected at 25 °C on a Bruker Avance NEO 600 MHz spectrometer equipped with a 5-mm BBO prodigy probe. Spectra were acquired using a pulse program in Bruker Topspin 4.1.4, processed and analyzed in Mnova 14.3.0. The IR spectra of purified PesA2(Q14K)-PesOHIC and PesA2(Q14K)-PesOHI core peptides were recorded with a PerkinElmer ATR-IR spectrometer.

**Site directed mutagenesis**

Plasmids encoding PesC mutants were constructed by polymerase chain reaction (PCR) based methods. Templates were amplified with Takara PrimeSTAR® Max DNA Polymerase Ver.2 and primers used are listed in Table S3. Successful mutations were confirmed by DNA sequencing.

***In vitro* assays**

For PesP1-P2 assays, the PesP1-P2 heterodimer (100 µM) was mixed with Ni-NTA-purified full length-PesOHIC peptide (100 µM) in reaction buffer (100 mM Tris, 150 mM NaCl, pH 8). The sample was supplemented with 1 mM $ZnSO_4$ and incubated at room temperature for 24 h. Then the sample was acidified prior to MS analysis.

For PesC assays, PesC (25 µM) was mixed with Ni-NTA-purified full length PesA2, PesA2-PesHI, or PesA2-PesOHI peptide (100 µM) in reaction buffer (100 mM Tris, 150 mM NaCl, pH 8). The sample was supplemented with 5 mM ATP, 20 mM $MgCl_2$, 10 mM L-glutamine and 1 mM TCEP. The reactions were incubated at room temperature, and after 30 mins, 1 or 6 h the mixture was acidified with formic acid (0.5% v/v final), desalted, and analyzed by MALDI-TOF MS or LC-MS. LC-MS analysis was performed on a Phenomenex Aeris 2.6 µm PEPTIDE XB-C18 LC column at 50 °C at a flow rate of 1 mL/min using 0.1% formic acid in $H_2O$ (solvent A) and 0.1% formic acid in acetonitrile (solvent B) with the following gradient program: 0–2 min 5% B, 2–12 min 5–95% B, 12–17 min 95% B, 17–19 min 95–5% B. For reactions with PesA2-PesOHI 8mer or 14mer peptide, 100 µM PesC was used and the mixture was allowed to react at room temperature overnight. For mechanistic and mutagenesis experiment, the standard reaction components were replaced by ATP analogs, $NH_4Cl$, or PesC mutants at the same concentration, and the assays were carried out for 6 h. For assays with the amide intermediate, 50 µL of the standard PesC in vitro reaction was quenched at 30 min by addition of 100 µL acetonitrile. After centrifugation, the supernatant was concentrated, and buffer exchanged three times to remove any unreacted ATP. Then the amide containing solution was incubated with 25 µM PesC with or without 5 mM ATP, 20 mM $MgCl_2$, incubated at room temperature for 6 h before analysis by LC-MS.

For MNIO activity assays, PesHI/PesXI/PesH (100 µM) was mixed with Ni-NTA-purified full length PesA2 peptide (100 µM) in reaction buffer (100 mM Tris, 150 mM NaCl, pH 8). The sample was supplemented with 330 µM iron(II) sulfate and 1 mM ascorbate. In PesX inhibition assays, 300 µM PesX was added to the PesHI reaction. The reactions were incubated at room temperature overnight and LC-MS analysis was performed using a Vydac C4 column at 50 °C at a flow rate of 1 mL/min using 0.1% formic acid in $H_2O$ (solvent A) and 0.1% formic acid in acetonitrile (solvent B) with the following gradient program: 0–2 min 5% B, 2–12 min 5–95% B, 12–17 min 95% B, 17–19 min 95–5% B.

For MNIO partner competition assays, PesHI (200 µM) was mixed with PesX (200 µM), and the mixture was incubated at 4 °C overnight. Then the mixture was passed through 0.5 mL of Ni-NTA resin, and the flow through and elution fractions were analyzed by SDS-PAGE.

**Table S1.** $^{1}H$ and $^{13}C$ NMR assignments of the LysC-digested PesA2(Q14K)-PesHIC peptide in 90% $H_2O$ and 10% $D_2O$ at 25 °C.

| Number | AA | NH<br>C=O | αH | βH | γH | δH | Others |
|---|---|---|---|---|---|---|---|
| 1 | Leu | N/A<br>170.5 | 4.02<br>51.9 | 1.72<br>40.0 | 1.65<br>23.8 | 0.95;<br>22.0, 21.7 | |
| 2 | Leu | 8.67<br>174.2 | 4.46<br>52.5 | 1.65<br>39.9 | 1.65<br>23.8 | 0.92;<br>21.1, 20.9 | |
| 3 | Gly | 8.32<br>168.8 | 4.17,<br>4.02<br>41.7 | | | | |
| 4 | Pro | N/A<br>174.1 | 4.43<br>60.3 | 2.27, 1.90<br>29.5 | 2.00<br>24.3 | 3.62<br>47.1 | |
| 5 | Arg | 8.44<br>173.7 | 4.36<br>53.4 | 1.81<br>28.1 | 1.62<br>24.4 | 3.22<br>40.7 | 157.0 (Cζ)<br>7.34 (NH) |
| 6 | Val | 8.32<br>173.3 | 4.24<br>59.3 | 2.11<br>30.2 | 0.93, 0.91<br>18.4, 17.3 | | |
| 7 | Asp-OH | 8.12<br>170.8 | 4.79<br>56.1 | 4.46<br>71.2 | | | 175.8 (Cγ) |
| 8 | Asn(CN) | 8.76<br>N/A | 5.11<br>37.8 | 2.97<br>36.6 | | | 172.6 (Cγ)<br>117.9 (Cα)<br>7.71,7.03<br>(amide) |

**In $^{1}H$ NMR, water peaks were referenced at 4.79 ppm.*

**Table S2.** $^{1}H$ and $^{13}C$ NMR assignments of the LysC-digested PesA2(Q14K)-PesOHIC peptide in 90% $H_2O$, 10% $D_2O$ and 0.1 % $d_2$-FA at 25 °C.

| Number | AA | NH<br>C=O | αH | βH | γH | δH | Others |
|---|---|---|---|---|---|---|---|
| 1 | Leu | N/A<br>170.5 | 4.03<br>52.0 | 1.72<br>40.0 | 1.66<br>23.8 | 0.95;<br>21.7, 21.7 | |
| 2 | Leu | 8.67<br>174.4 | 4.47<br>52.4 | 1.66<br>39.9 | 1.66<br>24.2 | 0.91;<br>21.0, 21.0 | |
| 3 | Gly | 8.35<br>169.3 | 4.18, 4.07<br>41.5 | | | | |
| 4 | Pro-OH | N/A<br>171.3 | 4.36<br>68.2 | 4.43<br>73.2 | 2.15, 2.08<br>32.3 | 3.75<br>44.7 | |
| 5 | Arg | 8.62<br>173.7 | 4.37<br>53.6 | 1.83<br>28.1 | 1.63<br>24.4 | 3.22<br>40.7 | 157.0 (Cζ)<br>7.33 (NH) |
| 6 | Val | 8.35<br>173.2 | 4.24<br>59.4 | 2.11<br>30.2 | 0.93, 0.92<br>18.3, 17.3 | | |
| 7 | Asp-OH | 8.14<br>170.7 | 4.82<br>56.0 | 4.50<br>71.1 | | | 175.6 (Cγ) |
| 8 | Asn(CN) | 8.77<br>N/A | 5.11<br>37.8 | 2.97<br>36.5 | | | 172.7 (Cγ)<br>118.0 (Cα)<br>7.71,7.03 (amide) |

**In $^{1}H$ NMR, water peaks are referenced at 4.79 ppm.*

**Table S3.** List of primers used to generate PesC mutants.

| | |
|---|---|
| PesC D334A-F | CTTAGCATGGCCCCAACAATAGTGG |
| PesC D334A-R | TATTGTTGGGGCCATGCTAAGCCC |
| PesC Y340F-F | ATAGTGGCATTTTACGCCTTGGGTG |
| PesC Y340F-R | AAGGCGTAAAATGCCACTATTGTTGG |
| PesC E363A-F | GTCCGACGCGATATTCGGCAGTTAC |
| PesC E363A-R | CGAATATCGCGTCGGACCCCGTC |
| PesC Y368F-F | TCGGCAGTTTCGATTGGCTTCATTC |
| PesC Y368F-R | AAGCCAATCGAAACTGCCGAATATC |
| PesC R443A-F | CGTTGCCTGCTGTGGACCTGGGG |
| PesC R443A-R | AGGTCCACAGCAGGCAACGCATCTG |

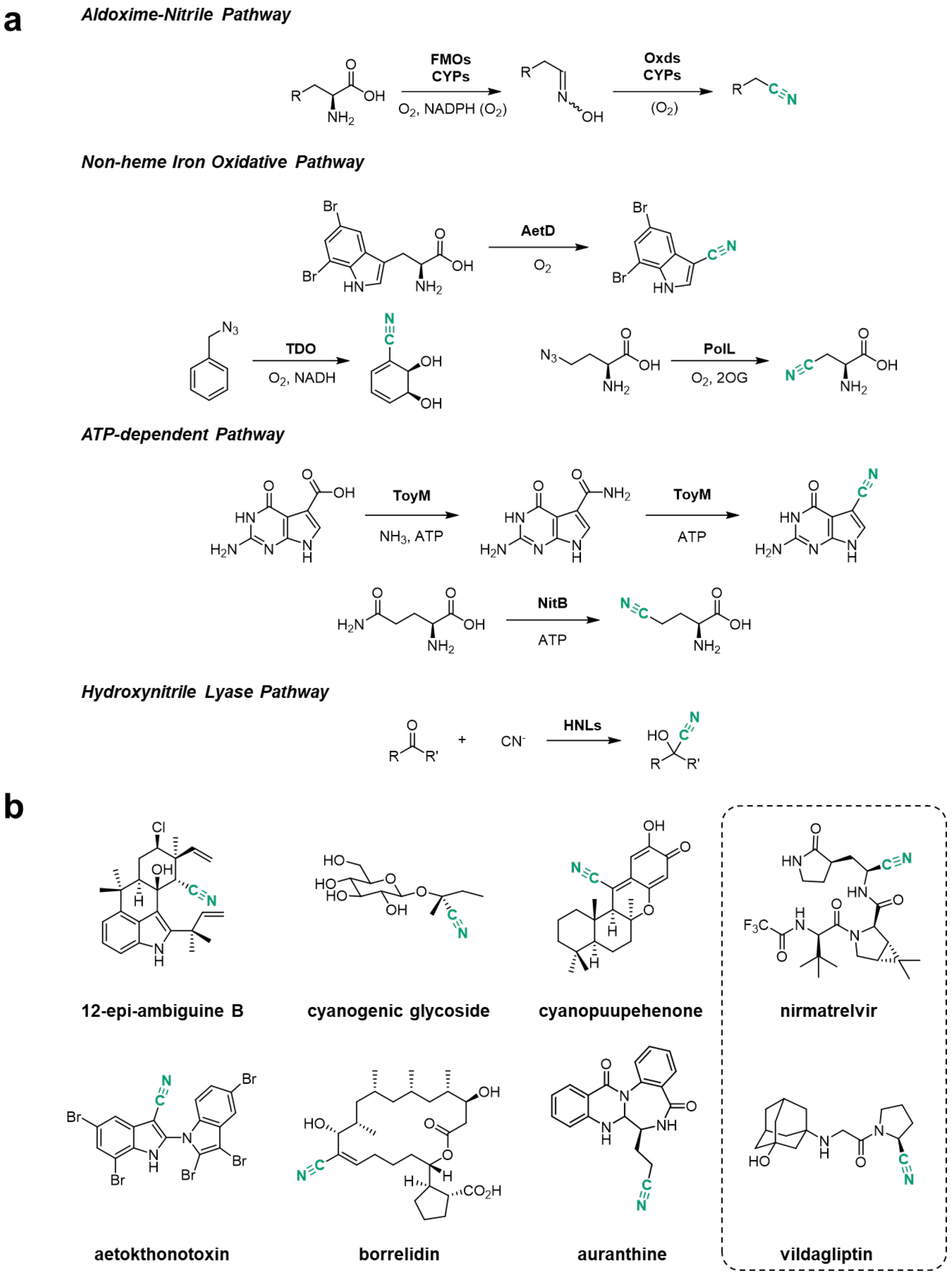


**Figure S1. a)** Schematic representation of known enzymatic nitrile formation pathways. In the aldoxime-nitrile pathway "Oxd" represents aldoxime dehydratase[3-6]; **b)** Structures of representative nitrile containing natural products; the structures of the nitrile containing drugs nirmatrelvir and vildagliptin are shown in dashed boxes.

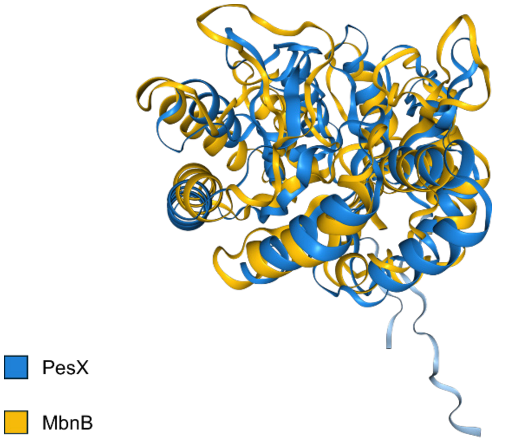


**Figure S2.** Overlay of the AlphaFold 3[7] generated PesX structure (shown in blue) and the crystal structure of MbnB (PDB ID: 7TCX, shown in yellow).

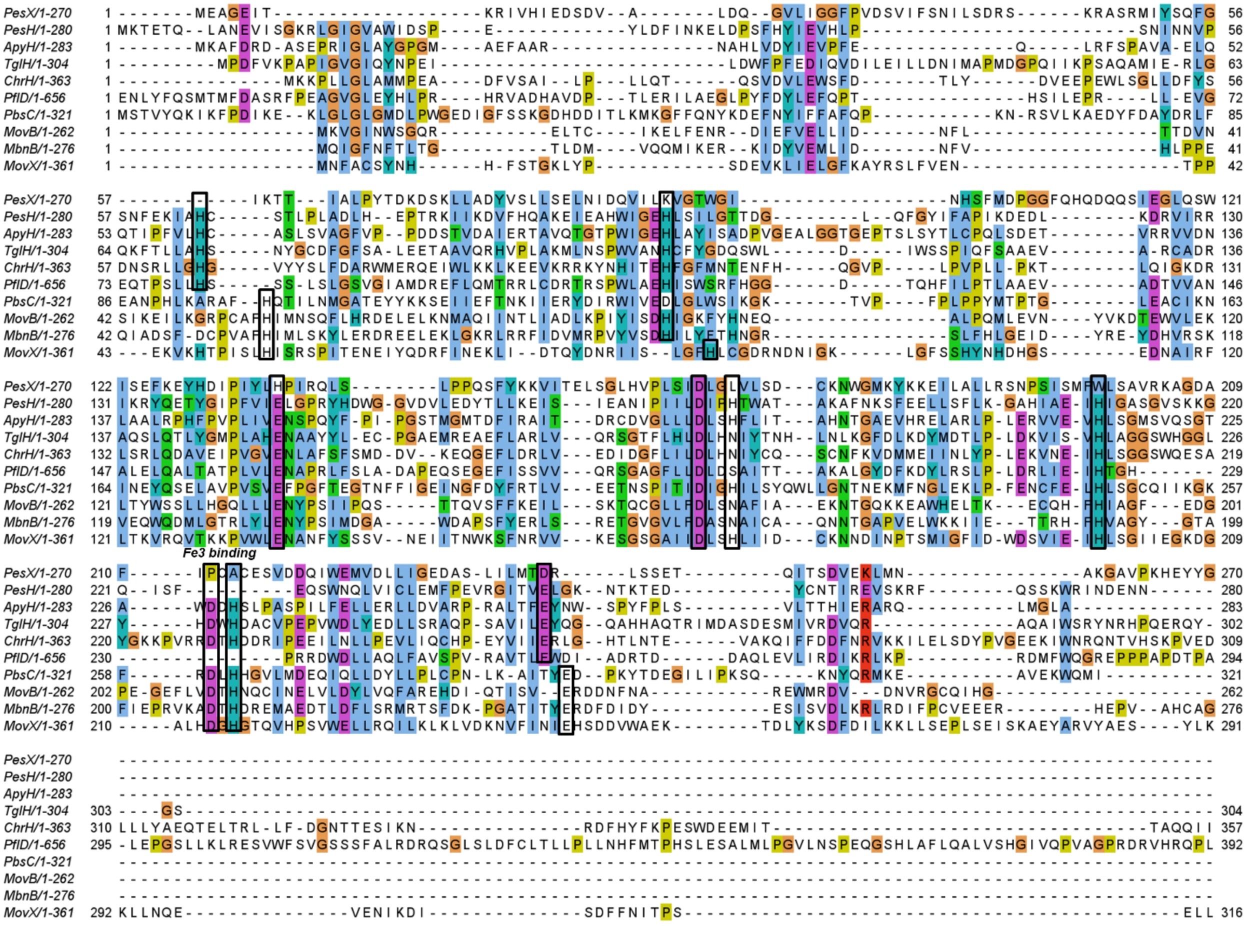


**Figure S3.** Multiple sequence alignment of representative characterized MNIO proteins with PesH and PesX. Conserved iron binding residues are labelled in black boxes. PesH and PflD lack ligands for Fe3 (see main text, also confirmed by AlphaFold modeling), whereas PesX lacks nearly all amino acid ligands to the iron ions that are in other MNIOs.

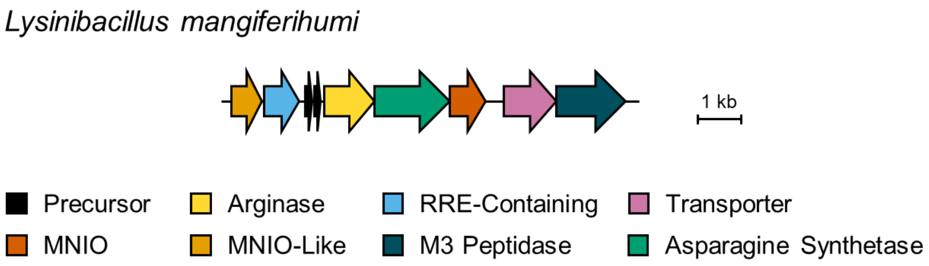


**Precursor1**: MMMDNKKIEEKLKELEEEITPNFFSESDVESEIPDSILGPLLRDRVDN
**Precursor2**: MDNKKVEEKLKELEEEITPNFFSESDVESEIPDSILGPLLRDRVDN

**Figure S4.** Composition and precursor peptide sequence of a representative *pes*-like BGC from *Lysinibacillus mangiferihumi*, where the gene encoding the PesO-like enzyme was absent and replaced by a gene encoding a putative arginase.

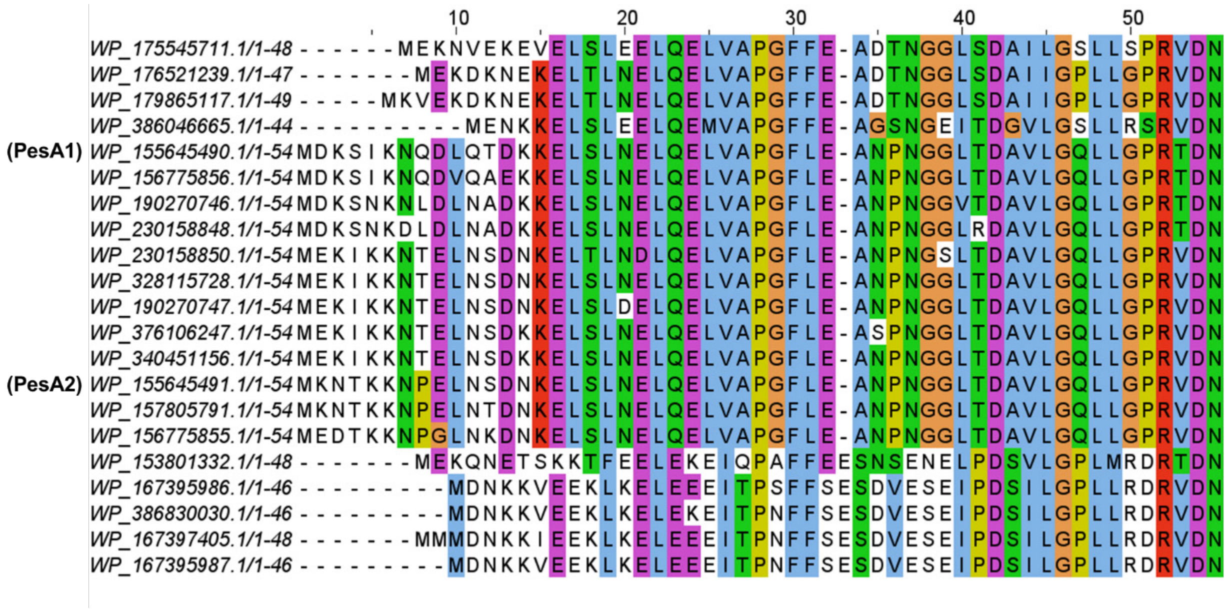


**Figure S5**. Multiple sequence alignment of 21 unique precursor peptides from the *pes* BGC, as well as orthologous clusters.

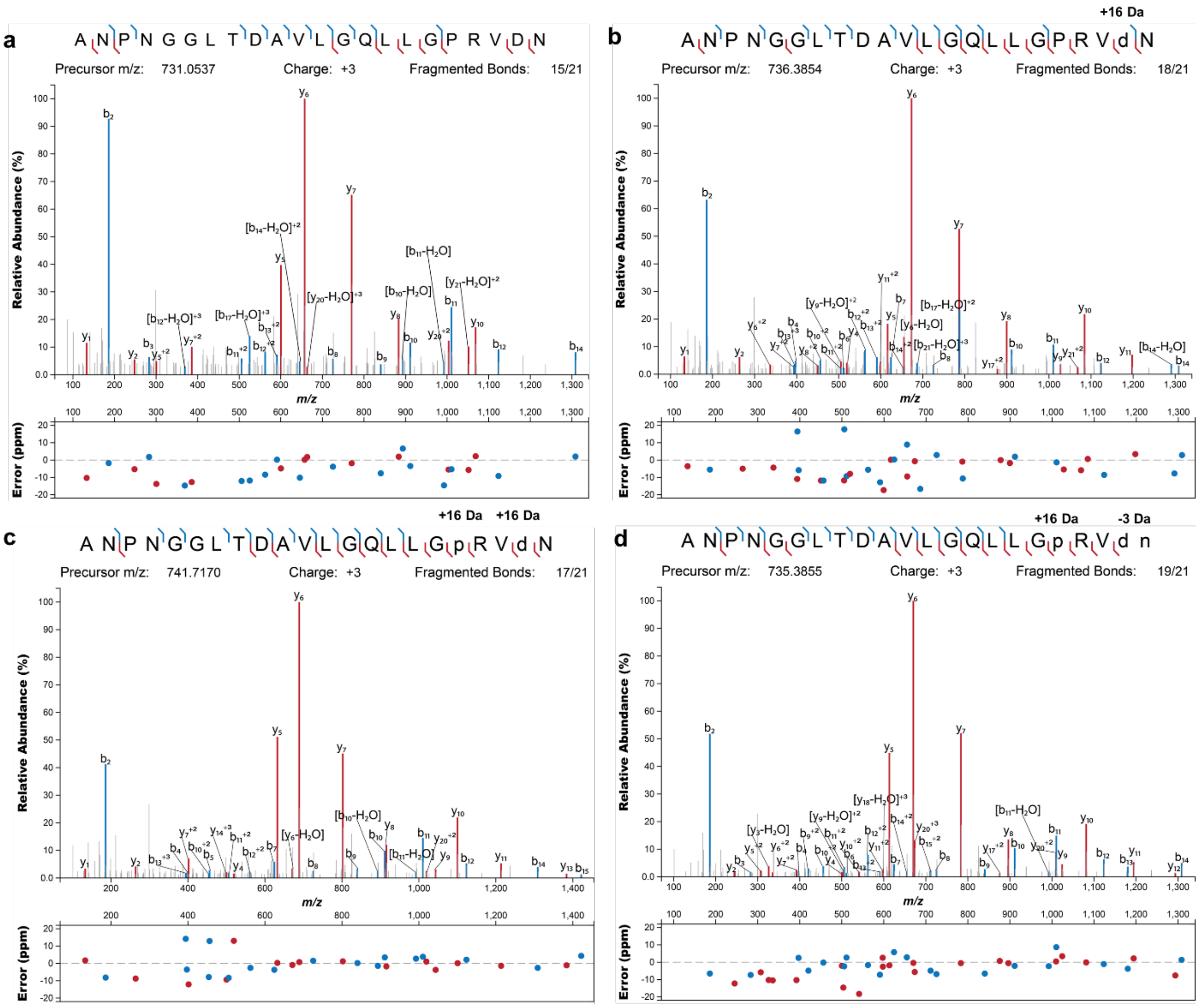


**Figure S6.** MS/MS analysis of the main species when PesA2 was co-expressed without or with modifying enzymes: **a)** PesA2, **b)** PesA2-PesHI, **c)** PesA2-PesOHI, and **d)** PesA2-PesOHIC. Prior to analysis, the peptides were digested with endoproteinase GluC. Fragment ion annotation was performed using the interactive peptide spectral annotator[1] with residues indicated in lower case p and d entered as hydroxylated residues (M+16), and in lower case d n as two residues that were hydroxylated and nitrile containing (+16 and –19 Da, respectively, to give a net change of −3 Da).

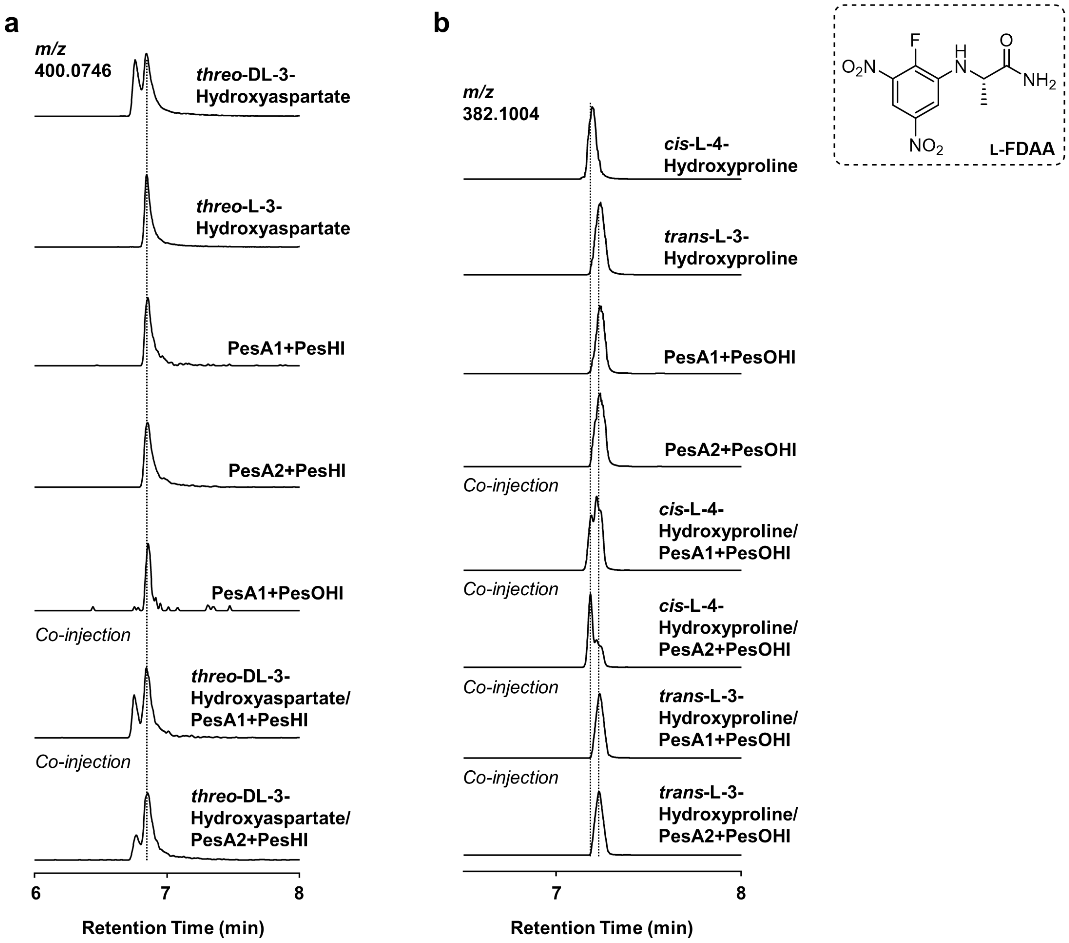


**Figure S7. a)** LC-MS analysis of L-FDAA (structure shown in dashed box) derivatized hydroxyaspartate from authentic standards or hydrolyzed PesA1-PesHI, PesA1-PesOHI, and PesA2-PesHI as well as co-injection of derivatized samples; **b)** LC-MS analysis of L-FDAA-derivatized hydroxyproline from authentic standards or hydrolyzed PesA1-PesOHI,and PesA2-PesOHI as well as co-injection of derivatized samples. Based on co-injection results, PesHI and PesO catalyze stereoselective C3-*S*-hydroxylation on Asp21 and Pro18 residues, respectively.

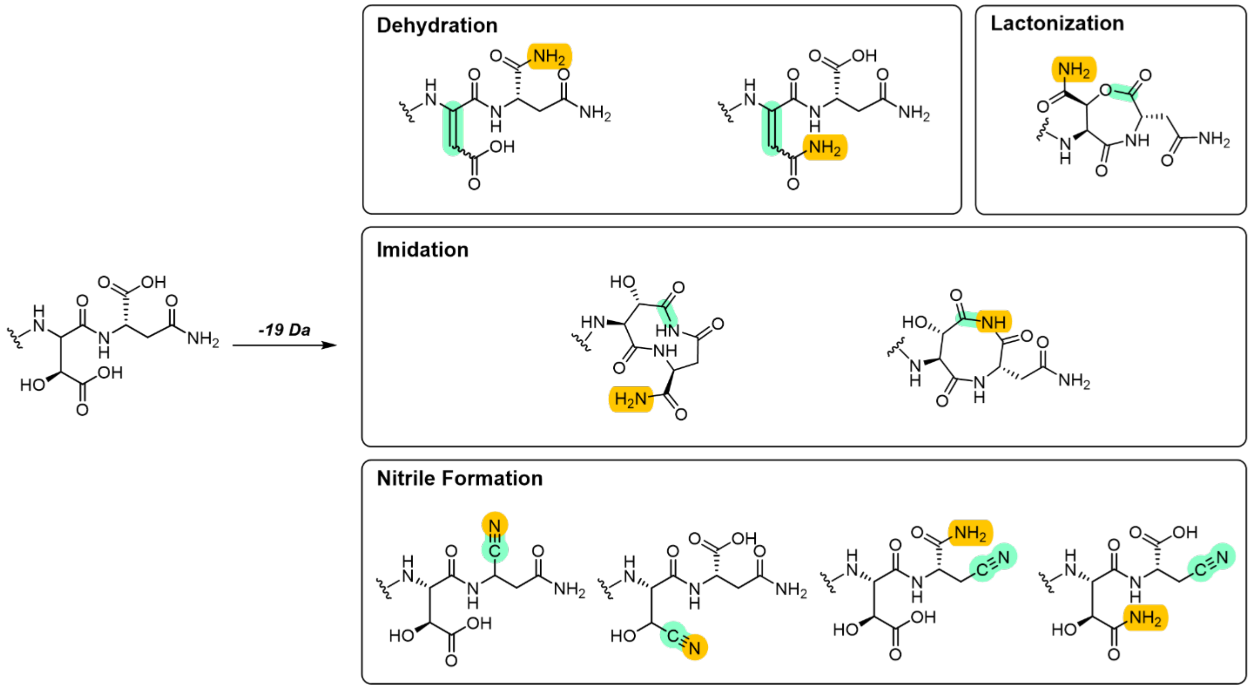


**Figure S8.** Proposed modifications at the precursor C-terminus that can account for the –19 Da mass shift after PesC modification. The amidation site is highlighted in yellow (–1 Da) and the proposed dehydration modification (–18 Da) is labeled in green.

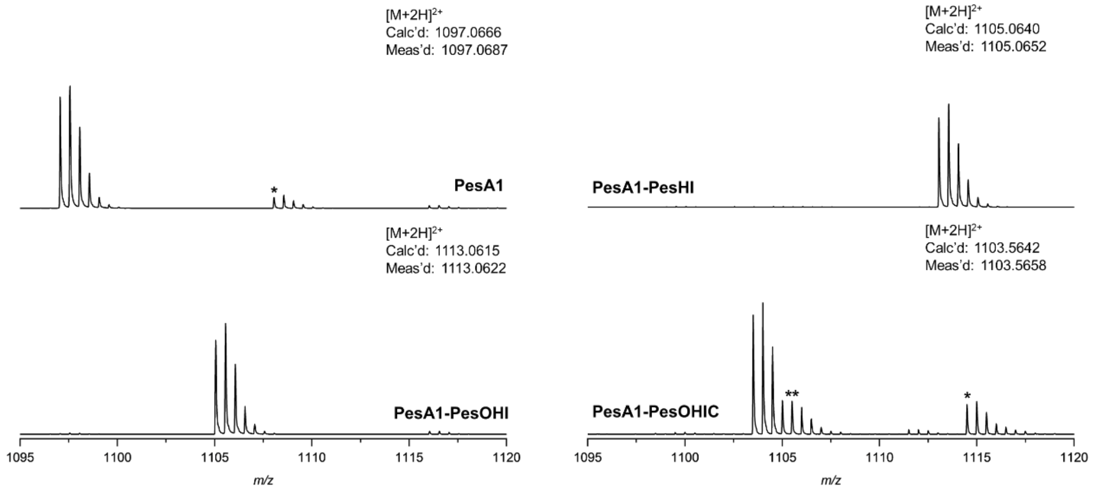


**Figure S9.** HR-MS analysis of unmodified precursor peptide PesA1 or PesA1 co-expressed with modifying enzymes. Prior to analysis, the peptides were digested with endoproteinase GluC. An asterisk (*) denotes [M+Na+H]$^{2+}$. In the PesA1-PesOHIC entry, the species marked with a double asterisk (**) is an unknown impurity from *E. coli* expression.

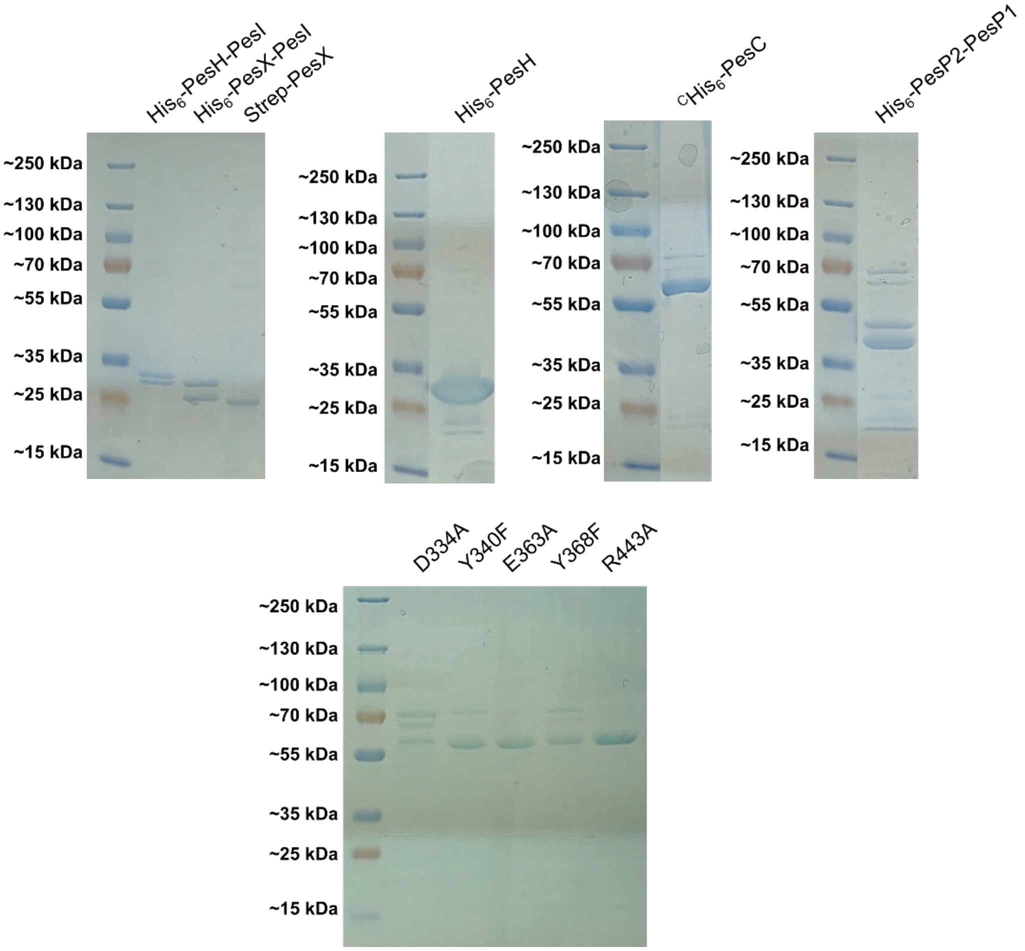


**Figure S10.** SDS-PAGE of proteins used in this study: $His_6$-PesH-PesI (32.1 kDa, 31.4 kDa), $His_6$-PesX-PesI (32.5 kDa, 31.4 kDa), Strep-PesX (31.8 kDa), $His_6$-PesH (32.1 kDa), $^{C}His_6$-PesC (68.4 kDa) and $His_6$-PesP2-PesP1 (48.2 kDa, 47.1 kDa) and $^{C}His_6$-PesC D334A, Y368F, Y340F, E363A, and R443A mutants.

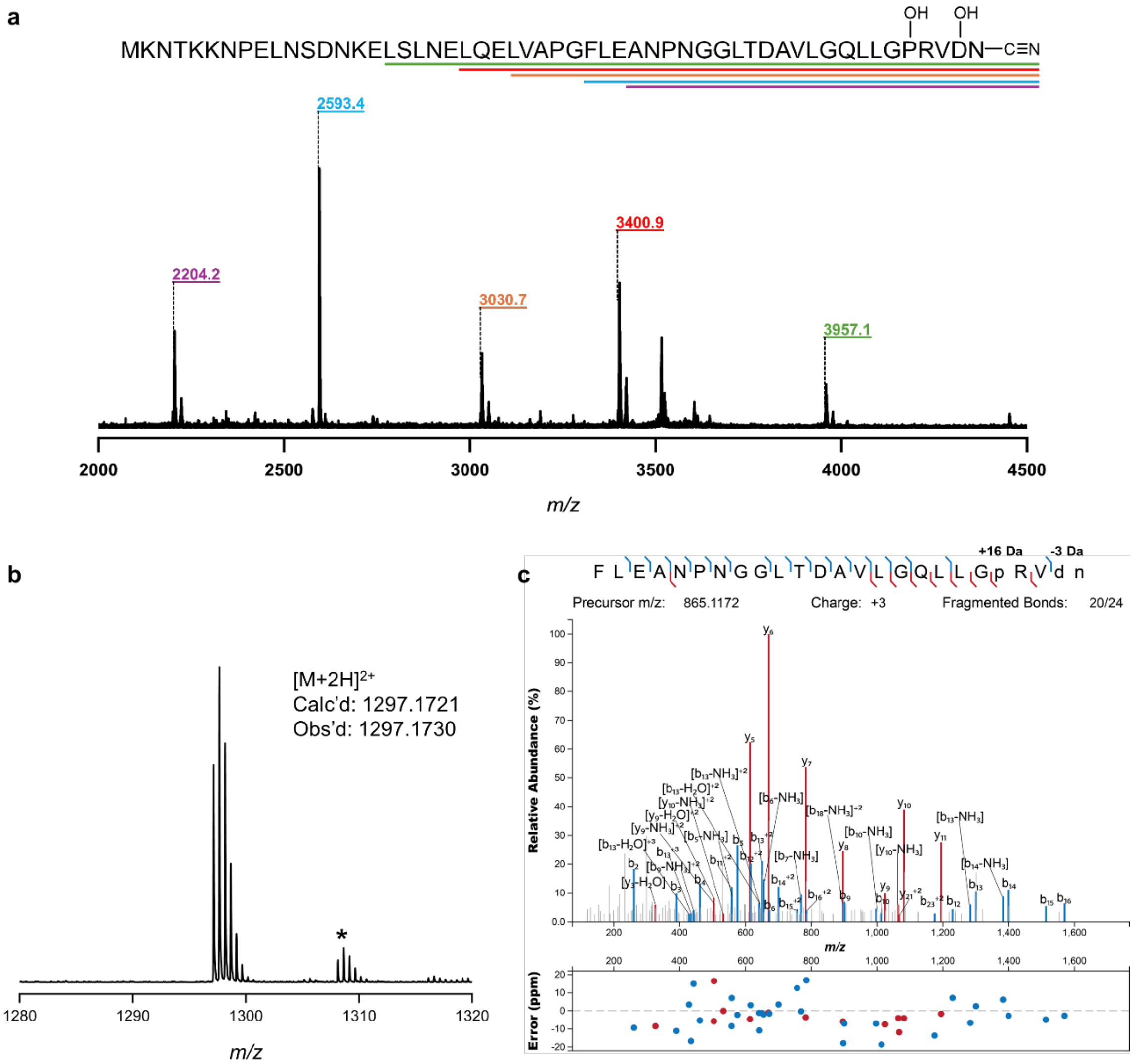


**Figure S11. a)** MALDI-TOF MS analysis of the full length PesA2-PesOHIC peptide after PesP2/P1 digestion. Peaks corresponding to the C-terminal fragment of the modified peptides are labelled, and the sequence of PesA2 is shown with colored lines indicating the sequence of each observed proteolytic product; **b)** HR-MS and **c)** MS/MS analysis of the major 25-mer core fragment of PesOHIC-modified PesA2 after PesP1/P2 digestion. An asterisk (*) denotes $[M+Na+H]^{2+}$. The fragment ion annotation was performed using the interactive peptide spectral annotator[1] with residues indicated in p entered as hydroxylated residues (+16 Da) and d n as two residues that were hydroxylated and containing a nitrile (net –3 Da).

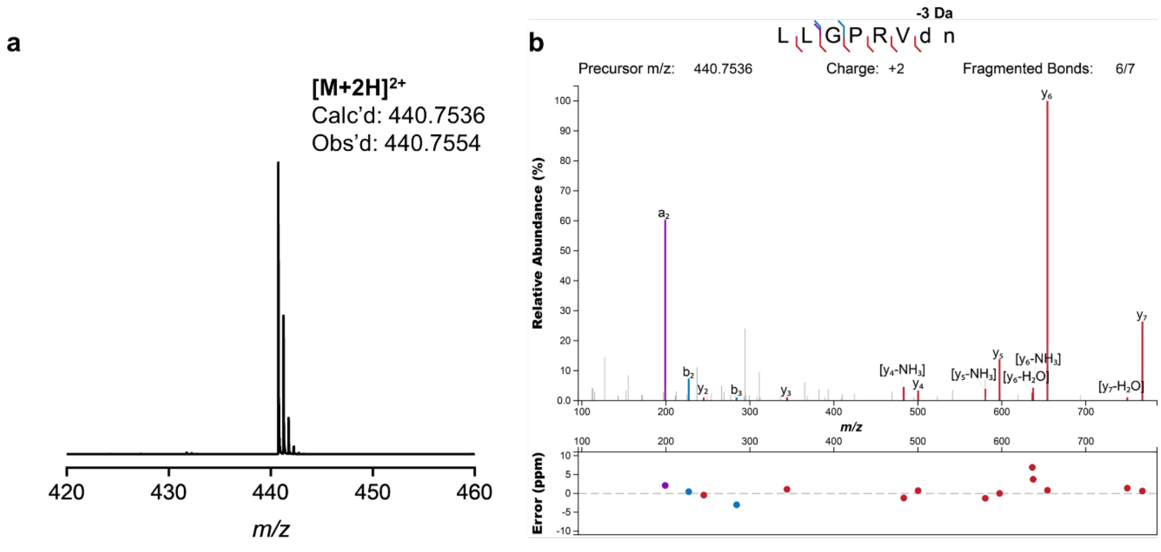


**Figure S12. a)** HR-MS and **b)** MS/MS analysis of the PesHIC-modified PesA2-Q14K mutant after LysC digestion and HPLC purification. Fragment ion annotation was performed using the interactive peptide spectral annotator[1] with residues indicated in d n as two residues that were hydroxylated and containing a nitrile (net –3 Da).

a

b

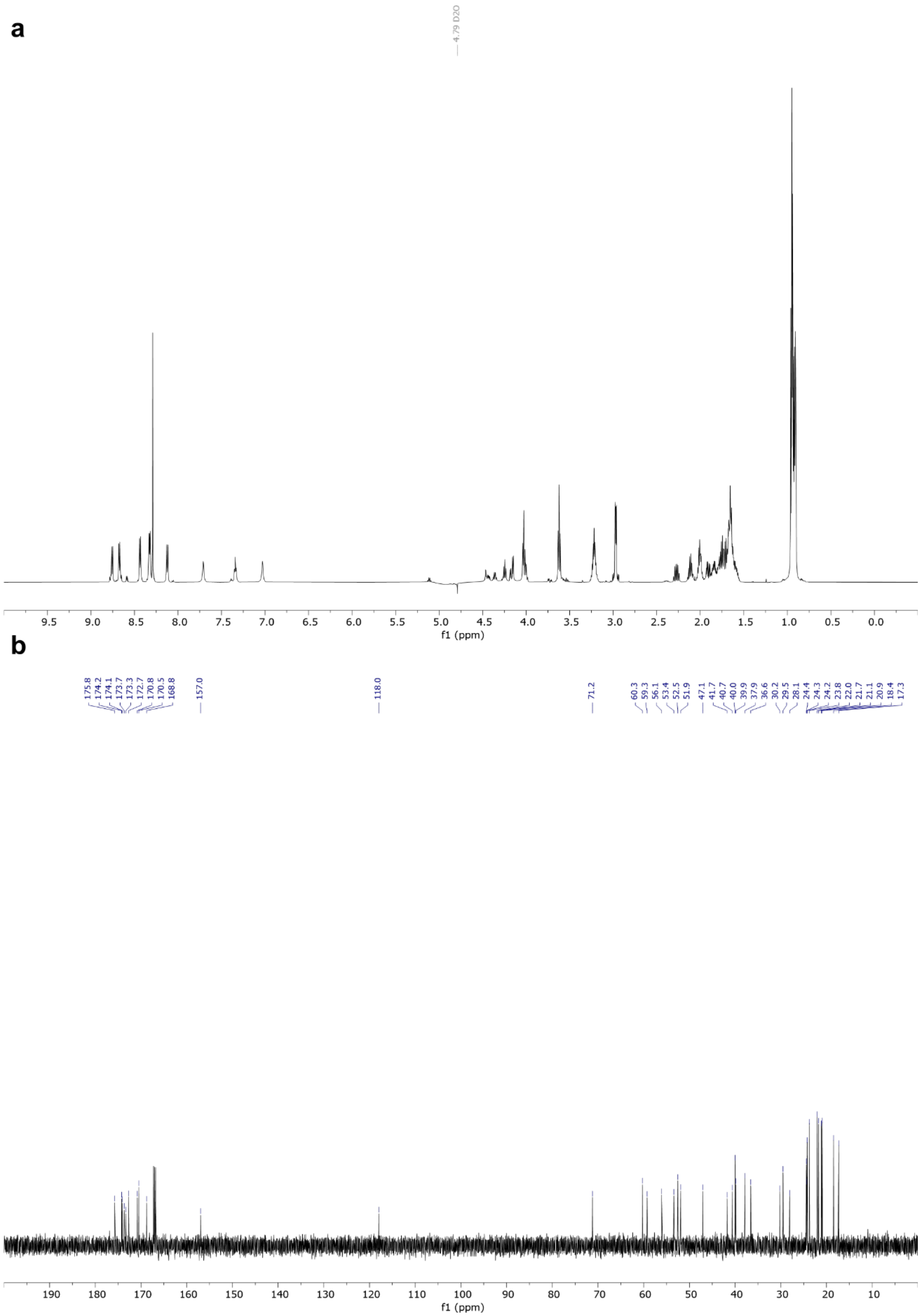

**Figure S13. a)** $^{1}H$ and **b)** $^{13}C$ NMR spectra of the PesA2-Q14K-PesHIC peptide after LysC digestion.

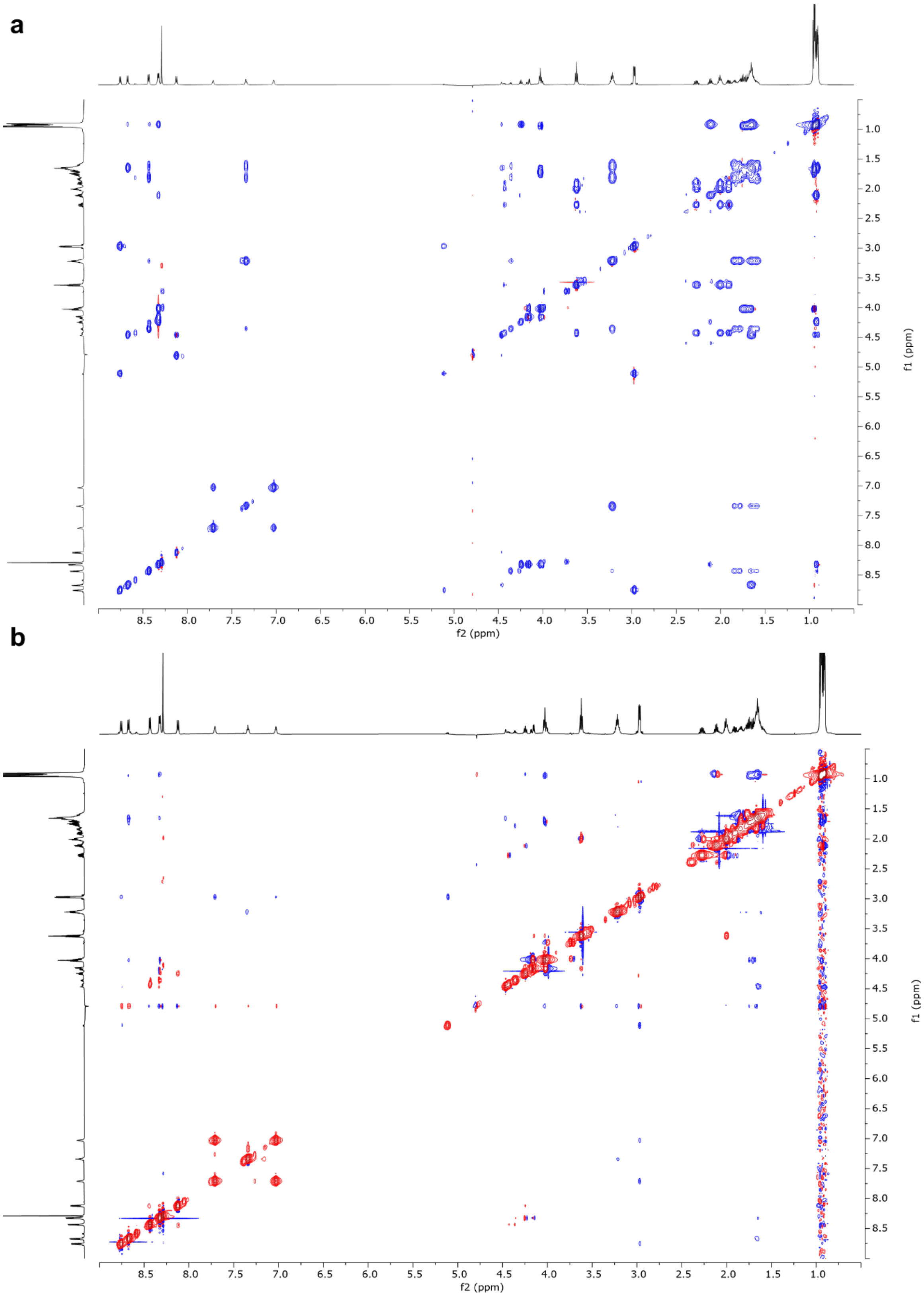


**Figure S14. a)** $^{1}H$-$^{1}H$ TOCSY and **b)** $^{1}H$-$^{1}H$ NOESY spectra of the PesA2-Q14K-PesHIC peptide after LysC digestion.

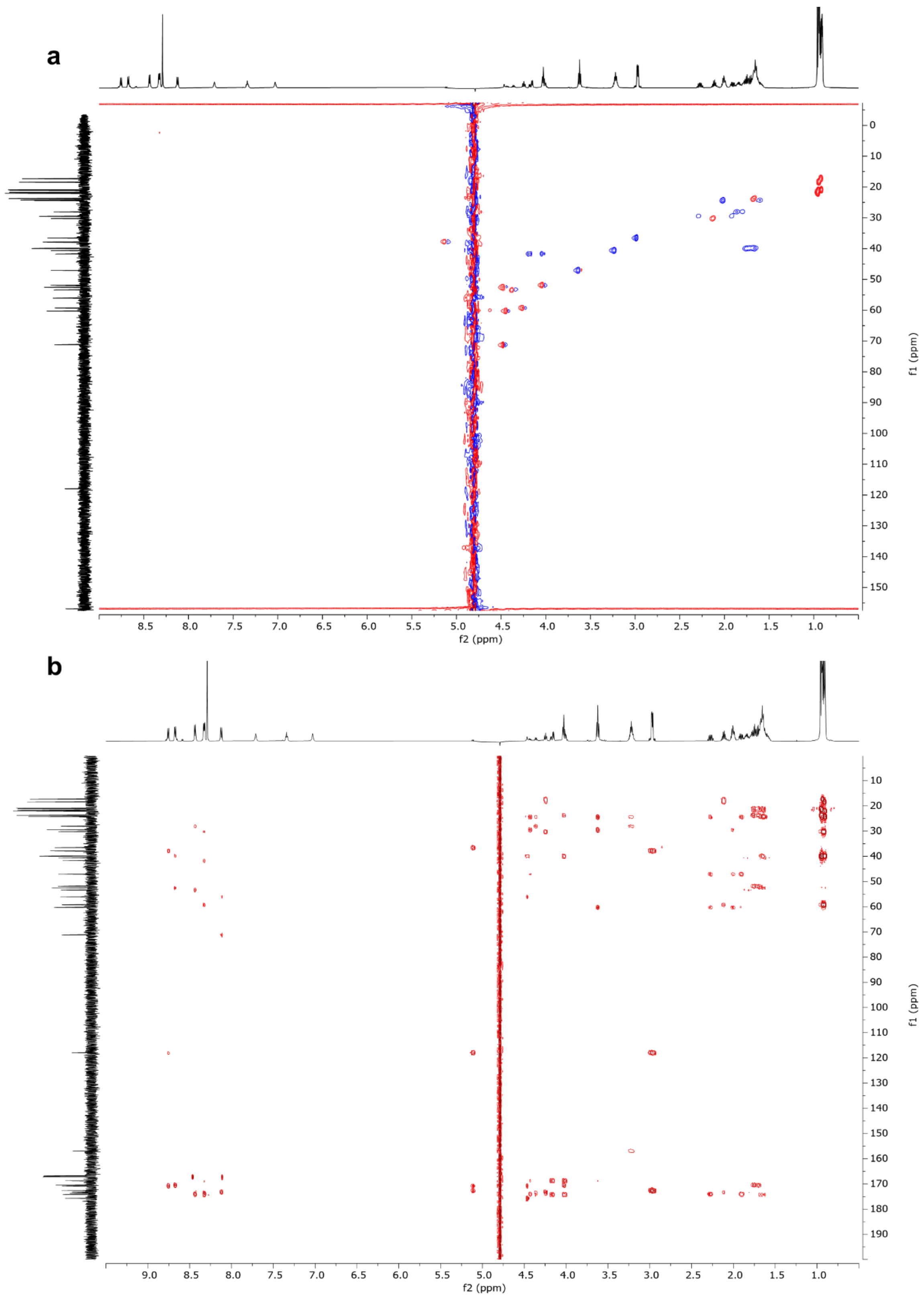


**Figure S15. a)** $^{1}H$-$^{13}C$ HSQC and **b)** $^{1}H$-$^{13}C$ HMBC spectra of the PesA2-Q14K-PesHIC peptide after LysC digestion.

a

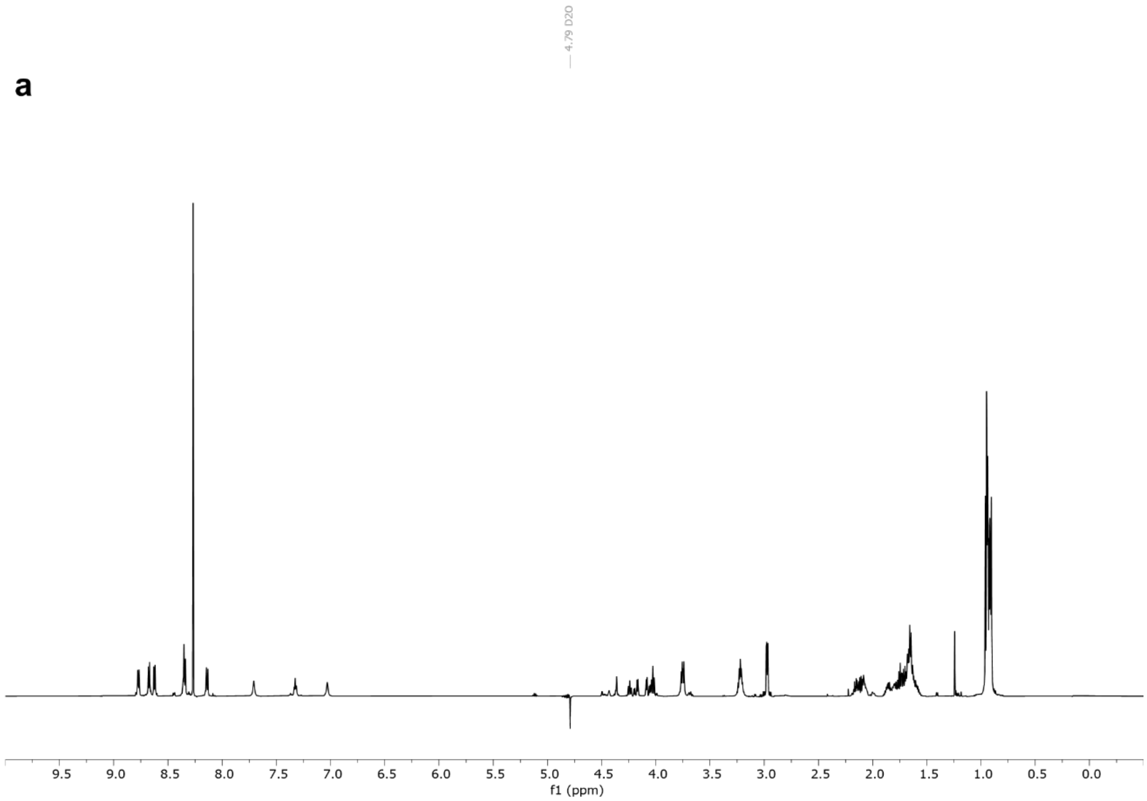


b

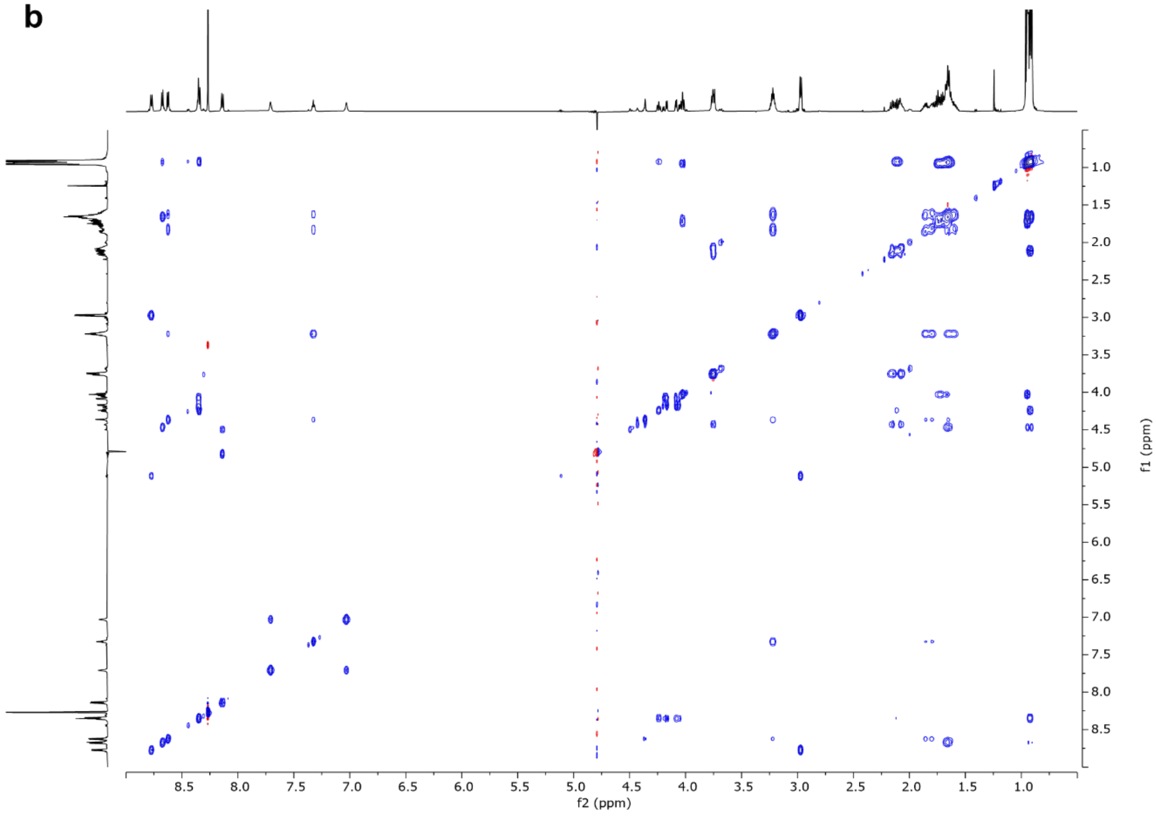


**Figure S16. a)** $^{1}H$ and **b)** $^{1}H$-$^{1}H$ TOCSY spectra of the PesA2-Q14K-PesOHIC peptide after LysC digestion.

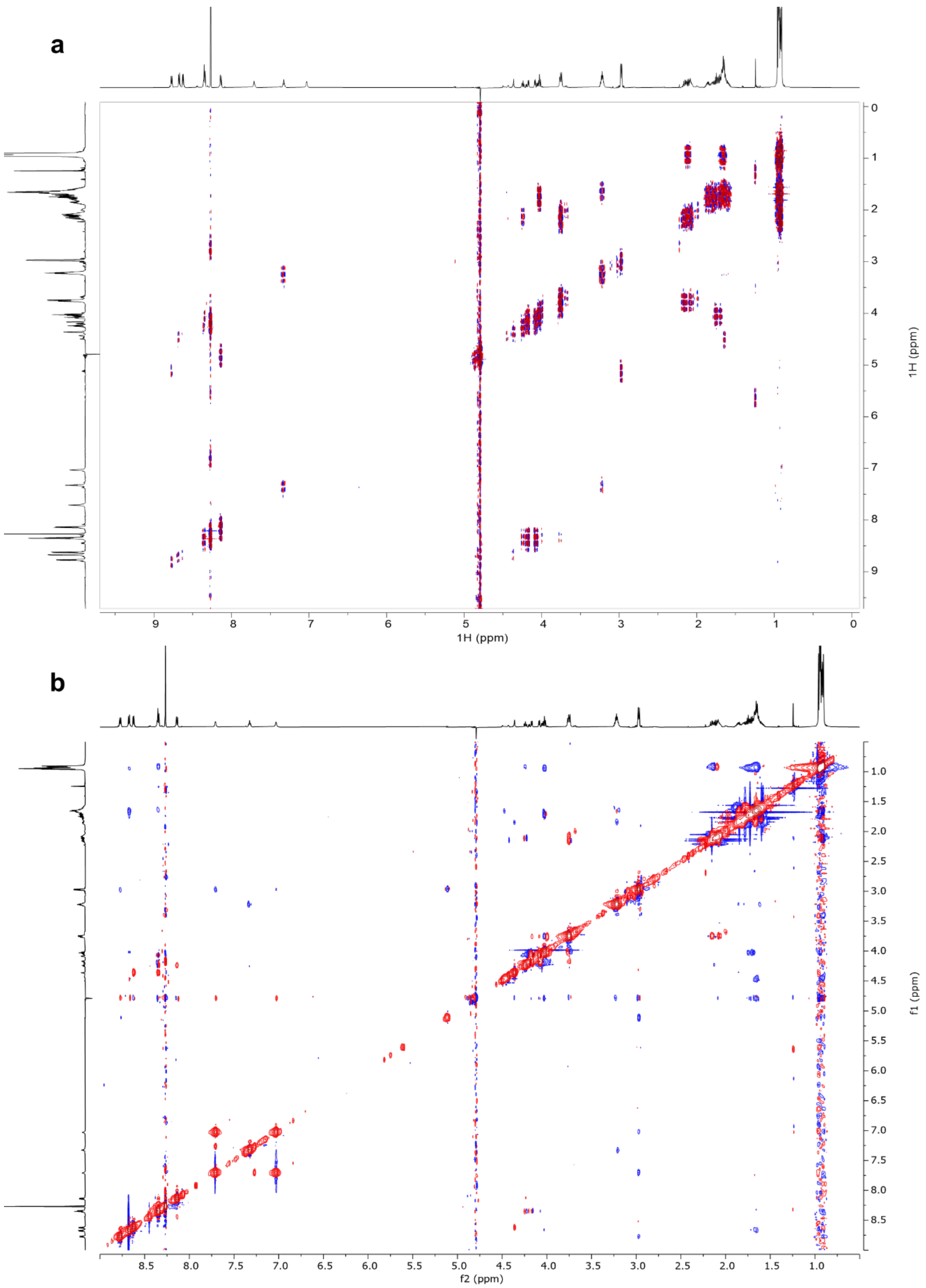


**Figure S17. a)** $^{1}H$-$^{1}H$ dqCOSY and **b)** $^{1}H$-$^{1}H$ NOESY spectra of the PesA2-Q14K-PesOHIC peptide after LysC digestion.

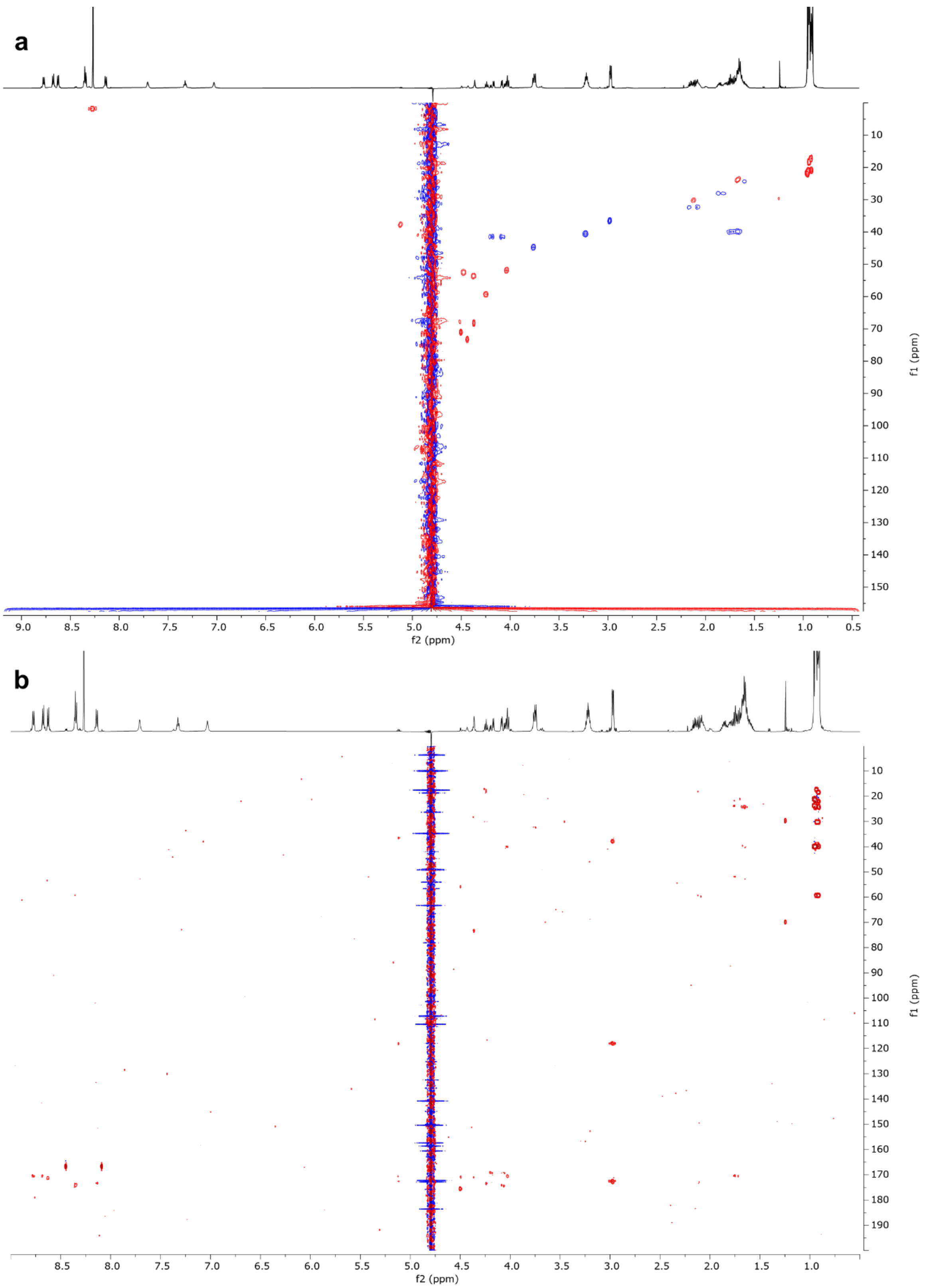


**Figure S18. a)** $^{1}H$-$^{13}C$ HSQC and **b)** $^{1}H$-$^{13}C$ HMBC spectra of the PesA2-Q14K-PesOHIC peptide after LysC digestion.

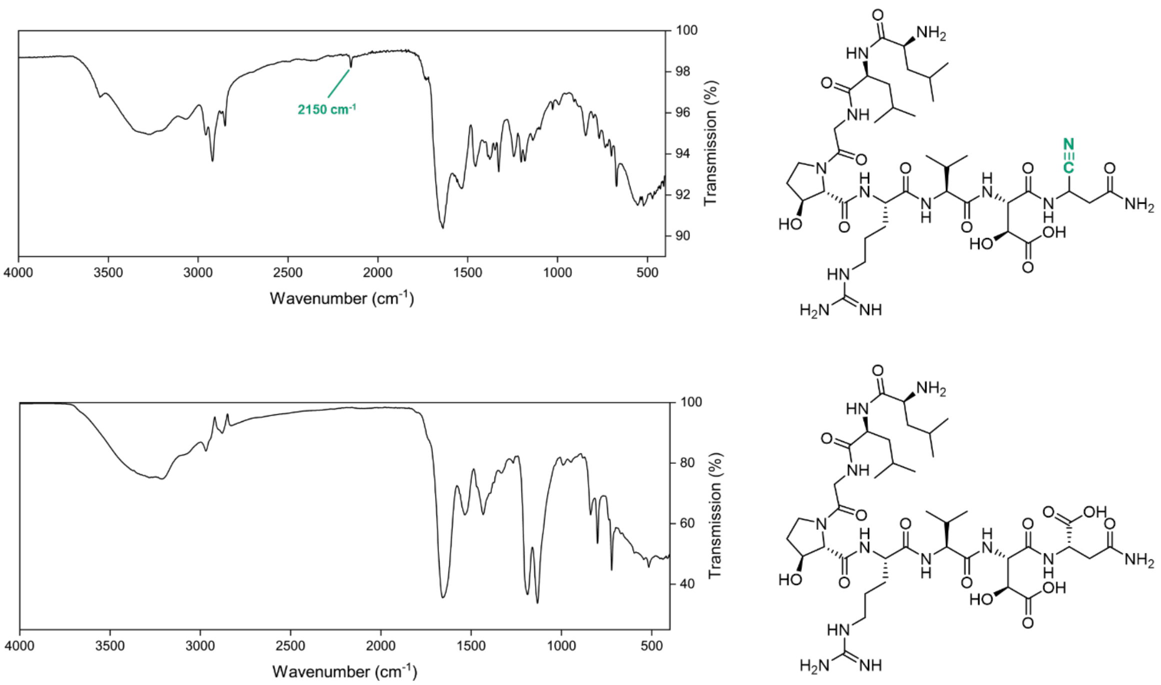


**Figure S19.** IR spectrum of the C-terminal 8-mer of PesA2-PesOHI (bottom) and PesA2-PesOHIC (top).

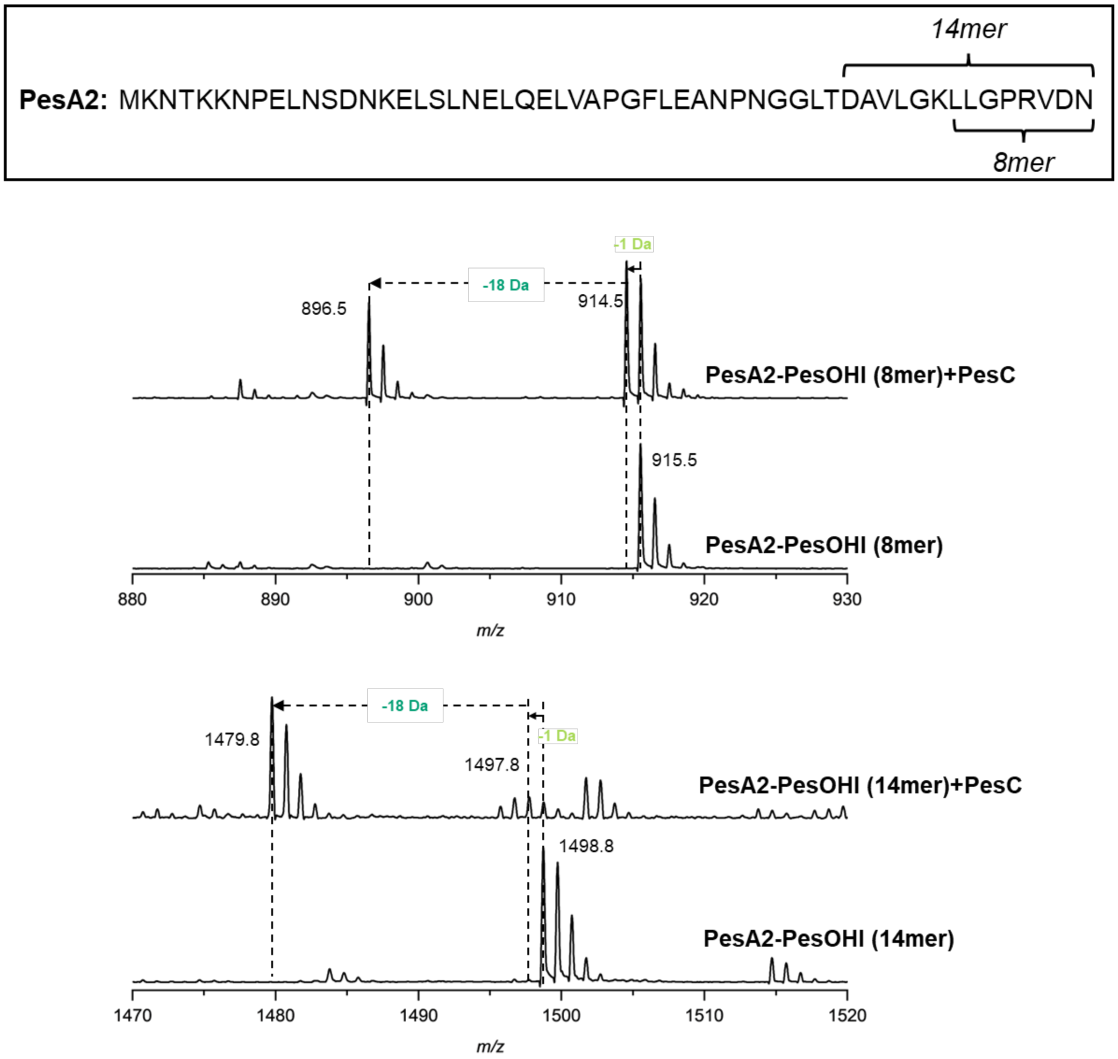


**Figure S20.** MALDI-TOF MS analysis of the *in vitro* PesC reaction with the C-terminal 8-mer or 14-mer of PesA2-PesOHI. The sequence of PesA2 as well as the sequences of the 8-mer and 14-mer are shown in the black box.

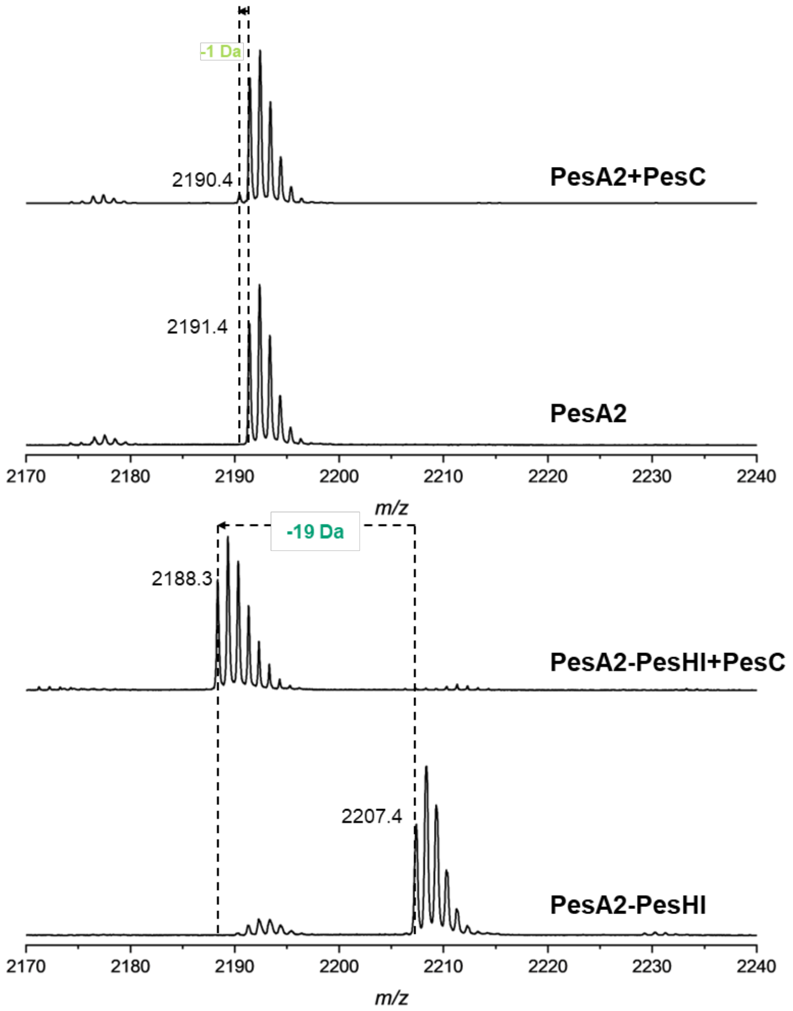


**Figure S21.** MALDI-TOF MS analysis of the *in vitro* PesC reaction with PesA2 or PesA2-PesHI after GluC digestion.

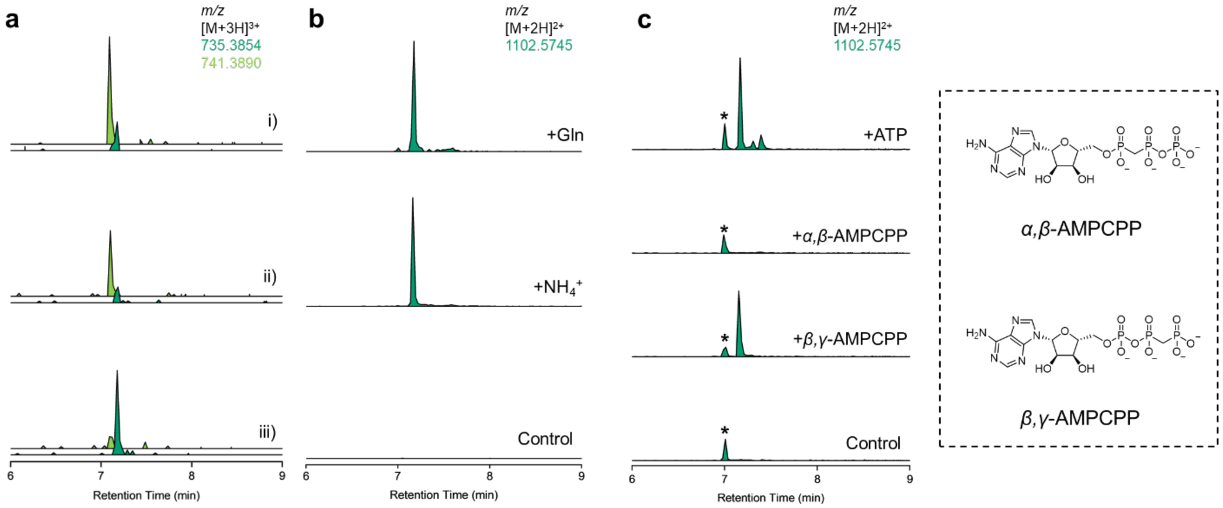


**Figure S22.** LC-MS analysis of PesC *in vitro* assays with PesA2-PesOHI after GluC digestion. **a)** The standard PesC reaction was first quenched at 30 min where amide (calculated mass $[M+3H]^{3+}$: 741.3890) is the major product over nitrile (calculated mass $[M+3H]^{3+}$: 735.3854). After protein precipitation by acetonitrile, concentration and buffer exchange, the amide containing solution was directly analyzed (i), or further incubated with PesC (ii), or PesC and ATP (iii); **b)** PesC assays with L-glutamine or $NH_4Cl$ as the nitrogen source, calculated mass for nitrile $[M+2H]^{2+}$: 1102.5745; **c)** PesC assays with ATP or the ATP analogs *α,β*-AMPCPP, and *β,γ*-AMPCPP; the structures of the ATP analogs are shown in the dashed box. An asterisk (*) denotes an unknown impurity during the analysis that is unrelated to the nitrilation reactivity of PesC and that is present in the negative control.

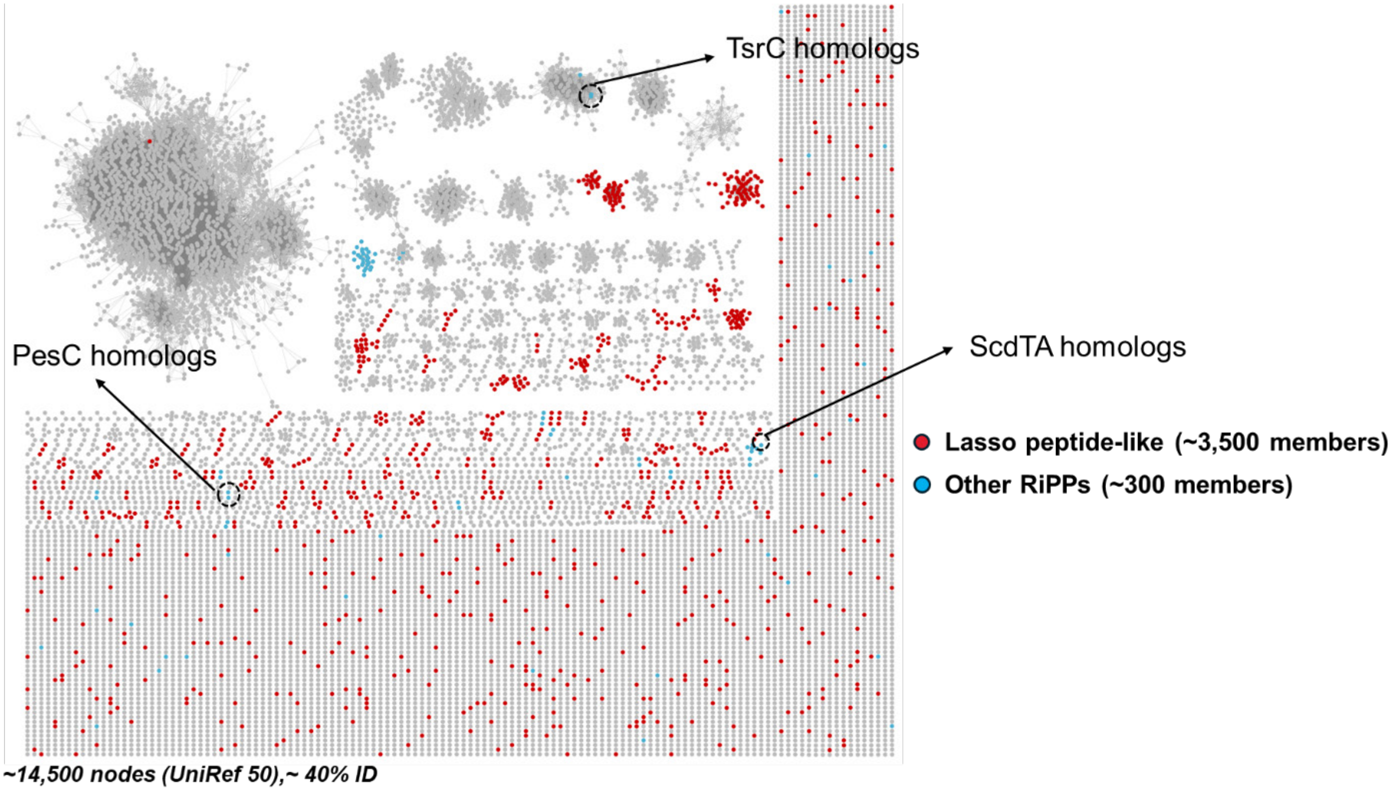


**Figure S23.** Sequence similarity network of the asparagine synthase protein family (PF00733) generated using the EFI-EST webtool[8] with the UniRef 50 database. The input accounts for ~14,500 entries as of November 2025, and the E-Value cutoff was set to 135 for members with ~40% identity. The genome neighborhood information of each entry was inspected manually with the assistance of Cytoscape and the EFI-GNT webtool. Members that co-occur with likely RiPP biosynthesis elements were picked. Predicted lasso cyclase-like members are labeled in red, with members involved in other potential RiPP biosynthetic pathways labeled in blue. Nodes representing characterized enzymes involved in C-terminal nitrilation (PesC) or amidation (ScdTA, TsrC) are labelled in dashed circles. The network is visualized and plotted with Cytoscape.

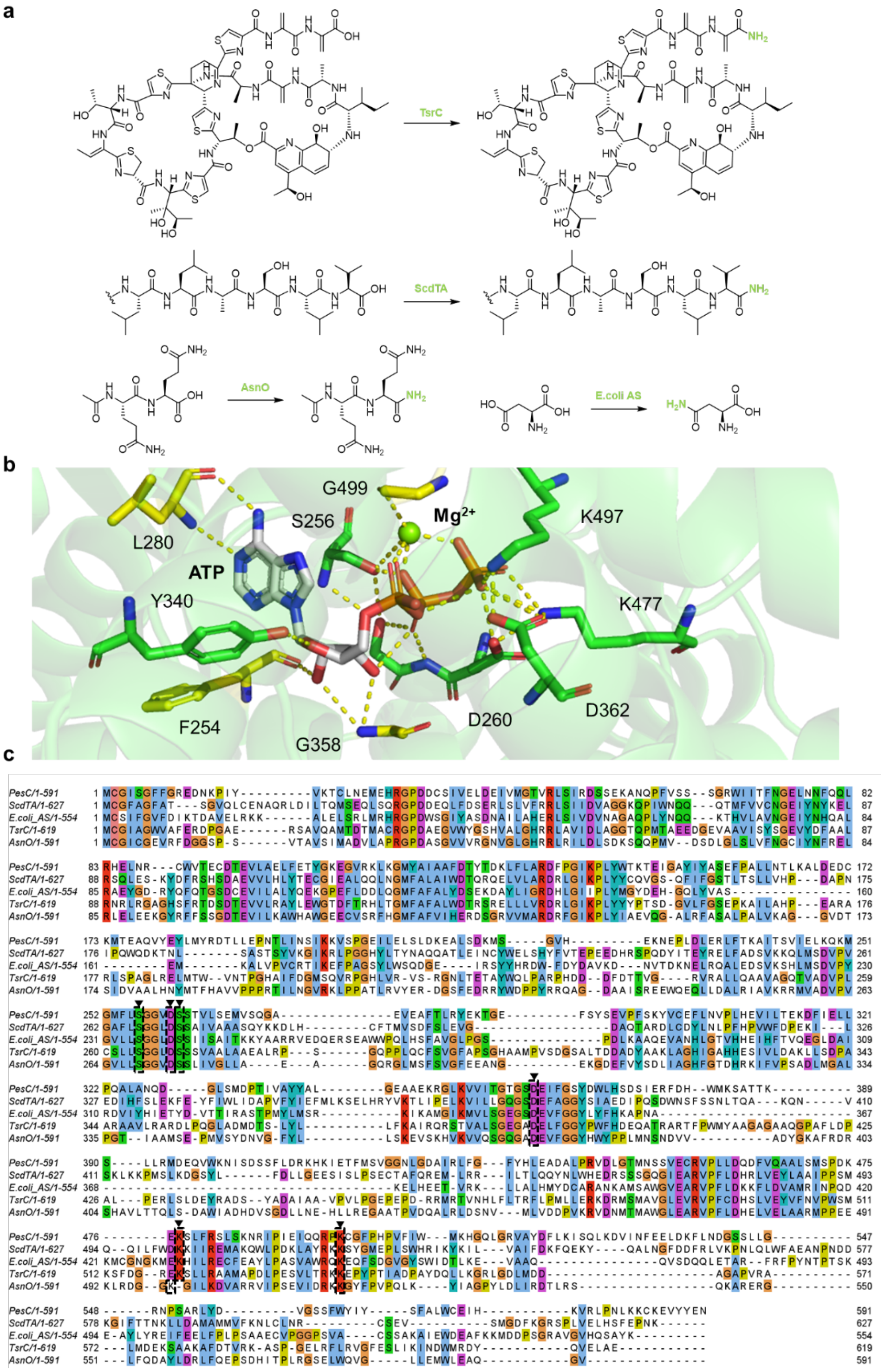


**Figure S24. a)** Amidation reactions catalyzed by AsnO, TsrC, ScdTA and *E. coli* AS; **b)** The ATP binding site of PesC in the AlphaFold 3 model. The residues involved in ATP and $Mg^{2+}$ binding are shown in green sticks (residues with backbone interaction to ATP and $Mg^{2+}$ are labelled in yellow sticks), and hydrogen bonds are indicated with yellow dashed lines; **c)** Sequence alignment of PesC, AsnO, TsrC, ScdTA and *E. coli* AS. Conserved residues that are predicted to be involved in ATP binding are labelled in dashed boxes.

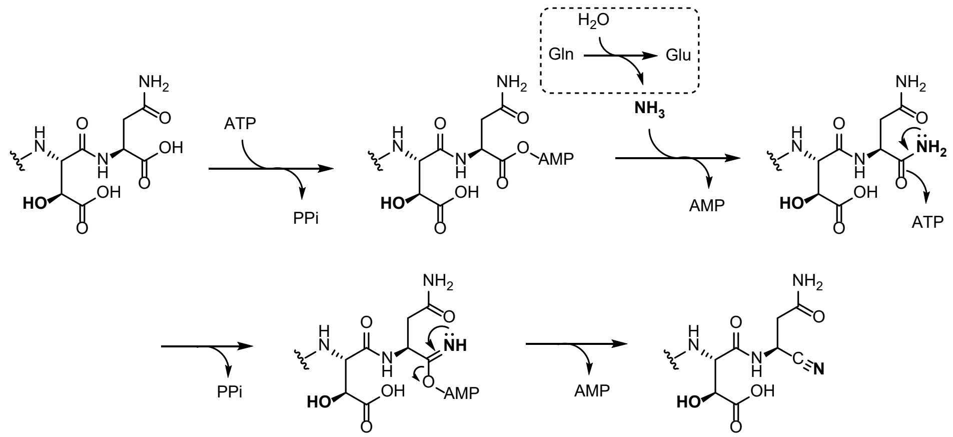


**Figure S25.** Proposed mechanism of PesC-catalyzed nitrilation.

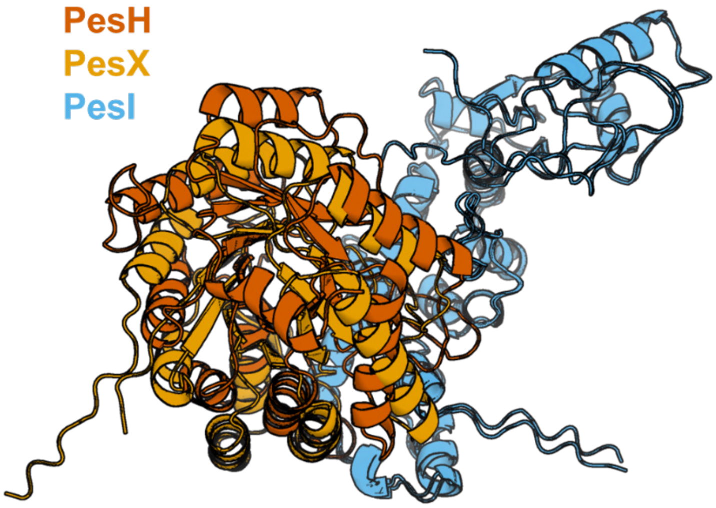


**Figure 26.** Overlayed AlphaFold 3 models of the PesHI and PesXI complexes.

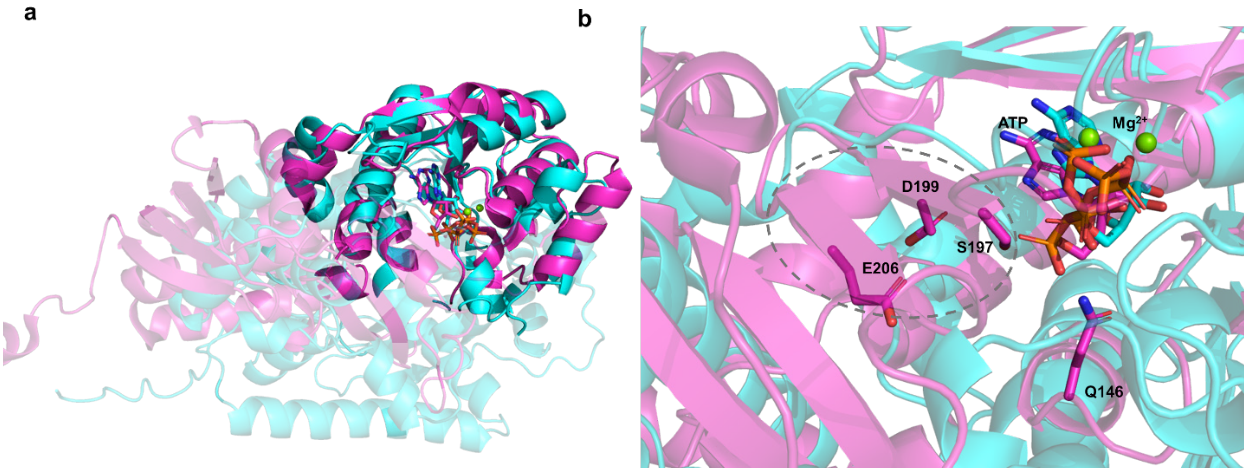


**Figure S27. a)** Superimposed AlphaFold 3 model of PesC (cyan) with the crystal structure of ArtA (PDB ID: 9LEH, magenta). The structural alignment reveals a similar ATP-binding domain; **b)** Zoom-in view of the PesC/ArtA active site, the key residues interacting with the substrate amide group in ArtA are labeled. These residues are not conserved in the active site of PesC. Additionally, the two β-strands that define the active site of ArtA[3] (marked in gray dashed circle) are not present in PesC, which likely adopts an exposed active site to allow large peptide substrate binding.

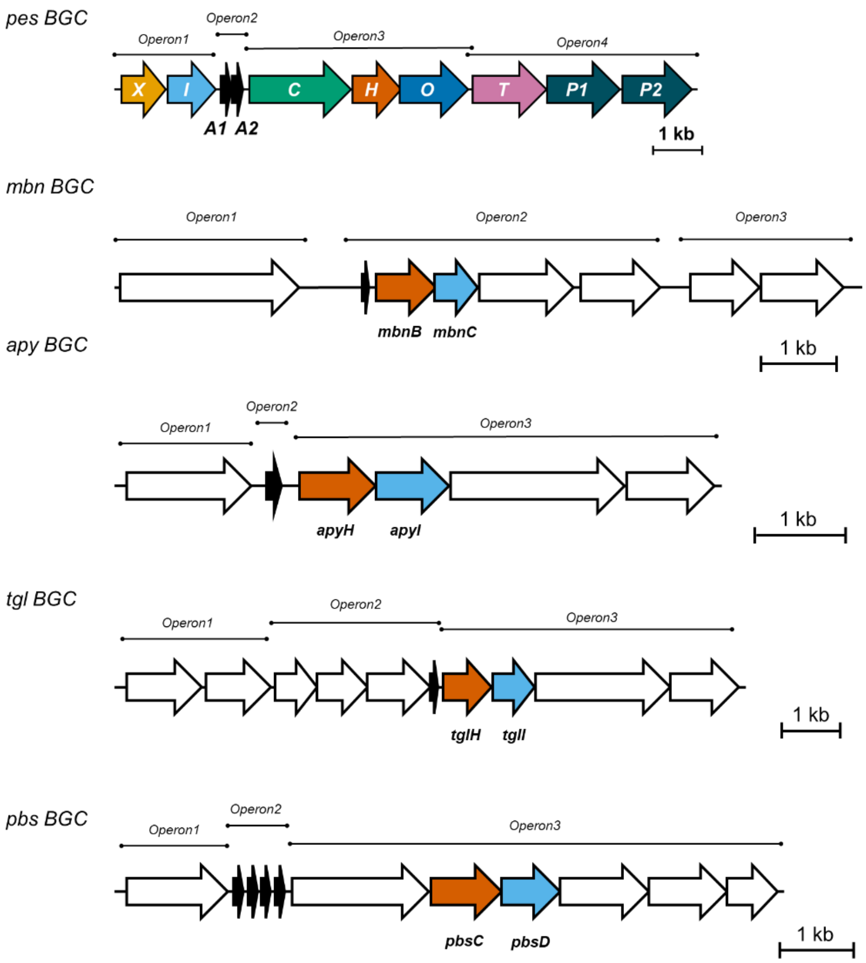


**Figure S28.** Operon predictions for the *pes* BGC and representative MNIO and MNIO partner protein encoding BGCs. The MNIO encoding genes are colored in red and the MNIO partner encoding genes are colored in blue. Predictions were done with Operon mapper[9].
